# Charting the Structure and Functional Landscape of the SF3B1 Intrinsically Disordered Region

**DOI:** 10.64898/2026.11.05.763689

**Authors:** S. Oliveti, K. Bera, L.M.R. Napolitano, M.A. Graewert, S. Di Marco, T. Staderini, B.P.O. Santos, L. Grisanti, S. Onesti, I.C. Felli, R. Pierattelli, A. Magistrato

## Abstract

Despite its essential role in pre-mRNA splicing, the structure and functional significance of the intrinsically disordered N-terminal domain (NTD) of SF3B1, a core component of the SF3b spliceosome complex, remain poorly understood. Here, we combine nuclear magnetic resonance (NMR) spectroscopy, small-angle X-ray scattering (SAXS), biochemistry, bioinformatics approaches, with molecular dynamics simulations to characterize the conformational landscape of the SF3B1-NTD and its phosphorylation-dependent remodelling. Specifically, we focus on the 186–341 region (^186–341^SF3B1), which contains at least four regulatory phosphorylation sites.

We reveal that ^186–341^SF3B1 is highly dynamic, lacking a stable secondary structure, and that phosphorylation of T211, T235, T313, and T328 induces a shift toward more compact states driven by the formation of intramolecular interactions. Additionally, we unveil that ^186–341^SF3B1 interacts with RNA primarily via the arginine residues flaking one of the conserved tryptophan-centred motifs present within SF3B1-NTD. These interactions are transient and heterogeneous, reflecting a nonspecific yet sequence-dependent recognition mechanism.

Collectively, this study provides a comprehensive description of the conformational landscape and functional features of the SF3B1-NTD, and expands the portfolio of its possible interactors, thus broadening our understanding of how intrinsic disorder, post-translational modifications, and sequence-encoded motifs cooperate to regulate pre-mRNA splicing.

**Highlights:** SAXS and NMR experiments reveal that SF3B1-NTD is intrinsically disordered

Phosphorylation compacts the SF3B1 intrinsically disordered region

SF3B1 intrinsically disordered region interacts with RNA at W200-centred motif

## Introduction

Intrinsically disordered proteins (IDPs) and proteins containing intrinsically disordered regions (IDRs) provide functional advantages that distinguish them from well-folded proteins. Lacking stable secondary and tertiary structures under physiological conditions, IDPs and IDRs can exploit their conformational flexibility to engage multiple binding partners with high specificity and relatively low affinity [1]. These flexible proteins are widespread across the proteome and are abundant among factors involved in pre-mRNA splicing [2]. Indeed, in the human spliceosome, approximately 40 % of proteins are predicted to be fully disordered and nearly 90 % contain at least one disordered segment [3].

The spliceosome catalyses the removal of introns from precursor messenger RNA (pre-mRNA) in the nucleus of eukaryotic cells, thereby contributing to produce mature protein-coding mRNAs and non-coding RNAs [4]. This machinery is composed of five small nuclear RNAs (snRNAs), each associated with a set of core proteins to form the U1, U2, U4, U5, and U6 small nuclear ribonucleoprotein (snRNP) particles, together with numerous additional proteins that bind and dissociate during the splicing cycle [5]. The spliceosome undergoes a series of conformational and compositional rearrangements, progressing through distinct intermediate states such as the E, A, B, Bact, B*, C, C*, P, ILS complexes. Spliceosome assembly is a highly dynamic process that is reinitiated for each splicing event [6].

Pre-mRNA splicing alterations are implicated in a wide range of human diseases such as cancer, neurodegeneration, and genetic diseases. Hence, to safeguard genome integrity, the spliceosome must ensure splicing fidelity at single nucleotide precision by accurately recognizing key signalling sequences [6,7]. Among them is the branch sequence (BS), which contains the branch-point adenosine (BPA), the nucleophile required for the first catalytic step of splicing [8].

BS recognition is mediated by the heptameric SF3b complex, a core component of the U2 snRNP. SF3B1 is the largest protein of SF3b complex, being composed by a long intrinsically disordered N-terminal domain (NTD; residues 1–463) and by a super helical C-terminal domain (CTD; residues 464–1304) composed of 22 HEAT repeats (Figure 1) [9]. SF3B1 is the most frequently mutated protein of the spliceosome and is also a therapeutic target for hematologic malignancies (i.e Myelodysplastic syndromes (MDS), Acute myeloid leukemia (AML), Chronic myelomonocytic leukemia (CMML))[10].

**Figure 1.**
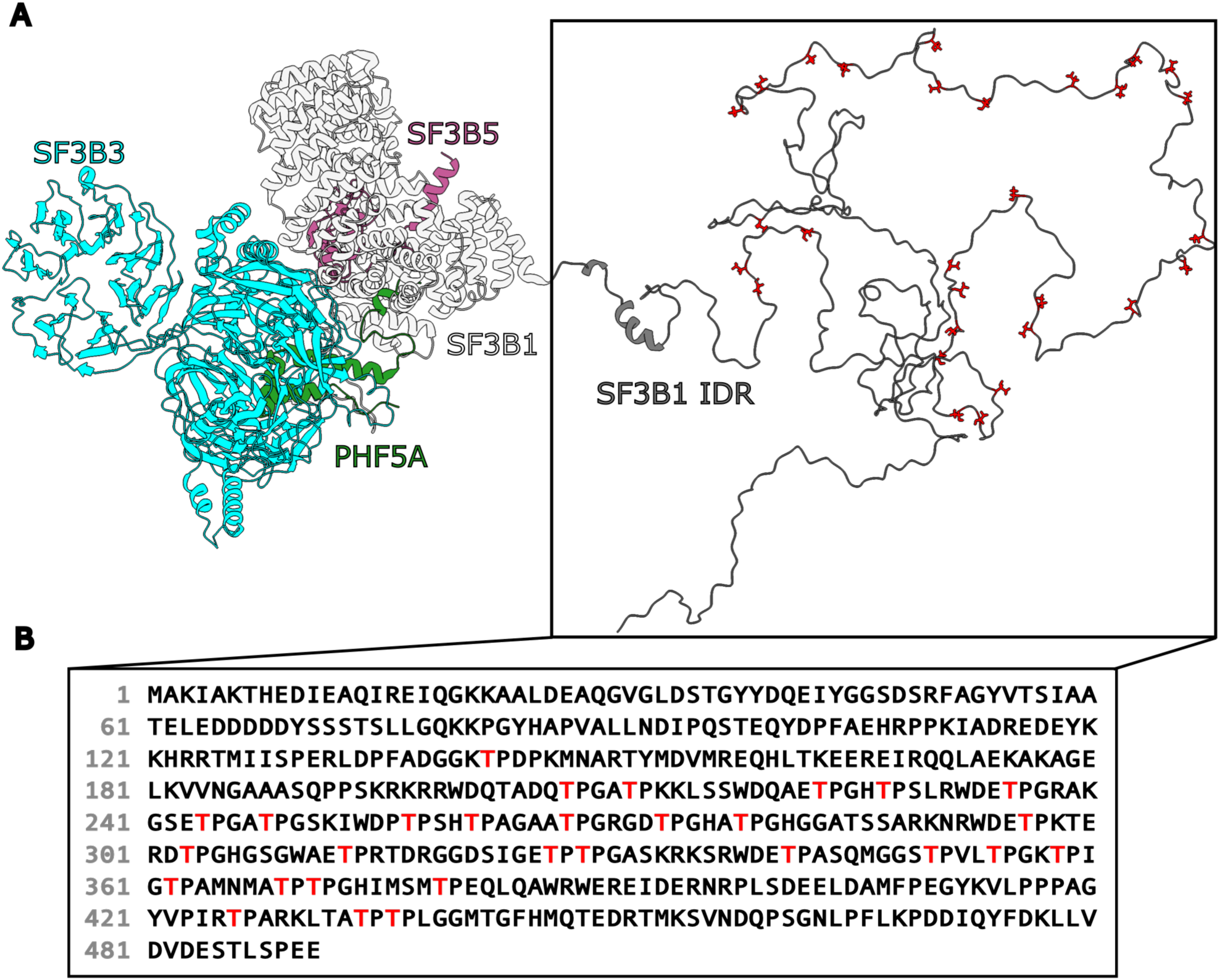
The SF3b complex. (A) Structure of the SF3b complex whose protein components SF3B3, SF3B5, PHF5A, and SF3B1 are shown in blue, green, pink, and white cartoon representations, respectively. The intrinsically disordered N-terminal region of SF3B1 is displayed on the right predicted with bAIes, which combines AlphaFold2 predictions with molecular modelling to generate atomic-resolution ensembles of intrinsically disordered proteins [11]. (b) Amino acid sequence of the intrinsically disordered N-terminal region of SF3B1. The ^186-341^SF3B1 fragment investigated in this study is shown in bold. In (a) and (B) phosphorylable threonine (T) and tryptophan (W) residues within the ^186-341^SF3B1 fragment are highlighted in red and blue, respectively.

While the function of C-term HEAT repeats is relatively well established, the role of the SF3B1 N-Term IDR remains elusive [12]. This region functions as a dynamic interaction hub for proteins and RNA and contains numerous threonine/proline (Thr/Pro) consensus sites required for phosphorylation by cyclin-dependent kinase 11 (CDK11) [13]. In particular, phosphorylation of the SF3B1 at T211, T235, T313, and T328 has been shown to trigger spliceosome activation, whereas a CDK11-selective inhibitor blocks the reaction prior to the first catalytic step [12,13]. These T/P consensus sites are often flanked by tryptophan residues preceded by nearby basic lysine or arginine residues and followed by aspartate or glutamate residues. This pattern is typical of many U2AF homology ligand motifs (ULMs) and is recognized by the several UHM (U2AF homology motif)-containing proteins [14]. Indeed, during early spliceosome assembly, particularly at the E-to-A complex transition, several UHM-containing splicing factors, including SF1, U2AF65, PUF60, Tat-SF1, RBM39, and SPF45 (RBM17), can bind the unphosphorylated ULM motifs of SF3B1 [14,15], regulating the recognition of the intron signalling sequences. At the spliceosome Bact complex, the Thr residues of the NTD-SF3B1 become phosphorylated and phosphorylation prompts the dissociation of ULMs from U2AF2 splicing factor [16]. This allows splicing to proceed until the formation of the C complex, where the second catalytic step occurs [17] and SF3B1-NTD is dephosphorylated [12].

Due to the key functional relevance of this SF3B1-IDR, and the sparse structural information currently available here we used an integrated experimental and computational approach based on nuclear magnetic resonance (NMR) spectroscopy, small-angle X-ray scattering (SAXS), bioinformatics, biochemistry, and molecular dynamics (MD) simulations to provide insights into SF3B1-NTD conformational ensembles and function [18]. We focused, in particular, on the 186–341 region (^186–341^SF3B1), which contains four threonine residues that undergo phosphorylation. This integrated approach revealed that the ^186–341^SF3B1 is completely disordered with only few hydrophobic interactions persistently established between the recurrent W residues and nearby hydrophobic residues. Conversely, MD simulations predict that actual phosphorylation of T211, T235, T313, and T328 residues shifts the conformational equilibrium of ^186–341^SF3B1 towards more compact structures. Since, IDRs are well known to interact with RNA [19], we also investigated the potential interactions of the SF3B1-NTD with different nucleic acid substrates. Binding assays revealed that also ^186–341^SF3B1 interacts with RNA, albeit with lower affinity than the full-length NTD construct (^1-463^SF3B1). Complementary NMR and MD simulations showed that the first of the seven ULM motifs, containing W200, established the strongest interactions with RNA, pinpointing the flanking arginine residues as the main mediators of these nonspecific interactions. Taken together, these findings identify the SF3B1-IDR as a regulatory hub whose interactome extends beyond protein partners, and more broadly, highlight the role of IDRs in regulating spliceosome function and plasticity.

## 2 Materials and Methods

### 2.1 Protein expression and purification

The pGEX-4T-3 SF3B1(1–463) plasmid was a gift from Dr. D. Blazek [13]. The protein was expressed in LB media in the *E. coli* pLysS BL21 (DE3) strain (Novagen). Cells were grown at 37°C to an OD_600_ of 0.7 and then transferred to 26°C. Recombinant protein expression was induced by the addition of 0.5 mM IPTG. After 20 h, cells were harvested by centrifugation. Cell pellets were thawed and resuspended in Lysis Buffer (50 mM Tris pH 7.5, 150 mM NaCl, 20% Glycerol, 0.5 mM EDTA, mM TCEP) supplemented with 1 mM PMSF, 15 μg/mL Benzamidin, 4 μg/mL Leupeptin and μg/mL Aprotinin. After cells disruption by sonicator using a KE76 probe (Sonoplus sonicator, Bandelin) the crude extract was centrifuged for 1 h at 30,000 *g* and the soluble fraction was incubated with 1 mL of Protino GST affinity resin (Macherey-Nagel). After 1 h of incubation at 4°C, the resin was washed with 20 resin volume of Washing Buffer (Lysis Buffer supplemented with 200 mM NaCl) and the protein was then eluted with the addition of 10 mM glutathione. As a final polishing step, the GST-tagged ^1-463^SF3B1 was loaded onto an analytical Superdex 200 Increase 10/300GL size exclusion column, using protein-storage buffer (50 mM Hepes pH 7.5, 150 mM NaCl, 0.5 mM EDTA, 1 mM TCEP). Proteins were finally flash frozen with liquid nitrogen and stored at −80°C.

The recombinant pET-28a(+) vector (Twist Bioscience), containing the ^186-341^SF3B1 gene, was transformed into *E. coli* BL21(DE3) competent cells for protein expression. For ^15^N and ^13^C-^15^N-labeled wild-type protein productions, a colony from a freshly transformed plate was inoculated in 100 mL of 2YT medium containing 50 mg L^-1^ kanamycin and incubated under shaking overnight at 37°C. Cells from the overnight cultures were collected by centrifugation and resuspended in 1 L of M9 minimal medium (supplemented with 1 mM MgSO_4_, 0.1 mM CaCl_2_, 1 mg L^-1^ biotin, 0.5 mg L^-1^ thiamine, 50 mg L^-1^ kanamycin, 3 g L^-1^ glucose/ ^13^C-glucose and 1 g L^-1^ ^15^NH_4_Cl) to give an initial OD_600_ of 0.15. Cells were incubated under shaking at 37°C. The culture was cooled down to 25°C when the OD_600_ reached a value of 0.5. The protein overexpression was induced with 1 mM isopropyl-β-D-thiogalactopyranoside (IPTG) at OD_600_ of 0.8 and cells were incubated overnight. The culture was harvested at 5000 rpm for 15 min and the pellet was resuspended in 50 mL of lysis buffer (50 mM Tris-HCl, 150 mM NaCl, 5 mM EDTA, 1 tablet of protease inhibitors, pH 7.2). Cells were disrupted by sonication on ice with cycles of 3 s with 12 s delay pulses for a total on-pulses of 3 min and centrifuged at 25,000 rpm for 25 min at 4°C. The soluble fraction was collected and diluted five times with a dilution buffer (50 mM Tris-HCl, 5 mM EDTA, pH 7.2) to reduce the salt concentration to 30 mM. The protein was purified by multi-step chromatography: one cationic exchange chromatography (CIEX), and a size-exclusion chromatography (SEC). All purification steps were carried out with an AKTA pure 25M (Ge HealthCare). The protein solution was loaded onto a 5 mL HiTrap FF SP column (Cytiva), previously equilibrated with binding buffer (50 mM Tris-HCl, 30 mM NaCl, 5 mM EDTA, pH 7.2). A linear gradient (0-100%) between binding buffer and elution buffer (50 mM Tris-HCl, 1 M NaCl, 5 mM EDTA, pH 7.2) was applied for 20 column volumes. ^186-341^SF3B1 was eluted with a NaCl concentration around 270 mM. Fractions were analyzed by SDS-PAGE, merged and concentrated to 5 mL using Amicon Ultra centrifugal filter units with a 3kDa MWCO (Millipore Sigma). The protein was heated up to 70-80°C for 20 minutes, rapidly cooled down in ice, centrifuged at 5800 rpm for 20 minutes and the supernatant was filtered. The protein was finally purified on a HiLoad 16/600 Superdex 75 pg column (Cytiva) with isocratic elution using the SEC buffer (50 mM sodium phosphate, 200 mM NaCl, 5 mM EDTA, pH 6.5). Fractions were checked by SDS-PAGE and stored at −80°C.

### 2.2 Oligonucleotide synthesis

All the oligonucleotides were chemically synthesized and purified by reverse-phase high pressure liquid chromatography (Biomers.net). Selected oligos were labelled with the fluorescent dye 6-Carboxyfluorescein (6-FAM). DNA and RNA oligos (Table S1) were resuspended in Tris–EDTA buffer (10 mM Tris–HCl pH 7.5, 1 mM EDTA) to a concentration of 100 μM and stored at −20°C. The RNase inhibitor RNase OUT (Thermo Fisher Scientific) was added to reactions containing RNA substrates (0.16 units/μL, 1:250).

For Electrophoretic Mobility Shift Assay (EMSA) assays, the substrates used were prepared using the following protocols: for *fork-DNA* (D1:D3) the fluorescent strand and the complementary strand were annealed at a 1:3 molar ratio in 20 mM Tris-HCl pH 8.0 and 100 mM NaCl. 100 μL reactions including 10 μM of each strand were heated to 95°C for 5 min and gradually cooled to 15°C at a rate of 1°C/min using a PCR thermal cycler (Eppendorf). For *D-loop* (D4:D9:D11) and *R-loop* (R4:D9:D11) the molar ratio used were 1:1.25:2.5 in 6 mM Tris–HCl pH 7.5, 7 mM MgCl_2_, 50 mM NaCl and 1 mM DTT. The annealing method used for D-loop and R-loop was heating at 99°C for 5 min followed by incubations at 67°C for 1 h, at 37°C for 30 min and at 25°C 3–4 h or overnight [20].Following electrophoresis in a native 10% acrylamide (29:1) TBE gel at RT in 1X TBE buffer, substrates were eluted from gel slices by dialysis in TE buffer.

For the NMR interaction experiments, a 20-nt RNA oligonucleotide (5′-CUACUACCCCCACCCUCACA-3′) obtained from Biomers.net and used without further purification, was employed.

### 2.3 Nucleic acid binding assays

Fluorescence anisotropy experiments were carried out in triplicate in a 384-well plate (Corning, 384-well, black, low volume, flat bottom, non-binding surface) at 25°C and fluorescence monitored by a microplate reader (Tecan Infinite F200 PRO) using a 495 nm excitation wavelength and a 525 nm emission wavelength. Every replicate contained a 30 μL solution containing 20 mM Tris-HCl pH 7.5, 50 mM KCl, 0.5 mM MgCl_2_, 10% Glycerol with 10 nM substrates and increasing concentrations (0– 1 µM) of the purified proteins.

For EMSA, binding reactions were performed in 20 mM Tris pH 7.5, 50 mM KCl, 0.5 mM MgCl_2_ and 10% Glycerol with 10 nM substrates and increasing concentrations (0–10 µM) of the purified protein in 20 μL of reaction volume, incubated at room temperature for 15 min. The reaction mixture was then loaded onto an 8% non-denaturing polyacrylamide gel and run at RT in TBE buffer. Fluorescent labelled substrates were detected by fluorescent scanner (ChemiDoc MP, BioRad) and quantification of protein bound nucleic acid was performed with Image Lab image software (BioRad). Each experiment has been repeated three times and plotted using Prism 8 (Graphpad).

### 2.4 SAXS data collection and analysis

Synchrotron SAXS data from the GST-^1–463^SF3B and ^186-341^SF3B1 fragments were collected on the EMBL P12 beamline [21] at PETRA III (DESY, Hamburg, Germany), using a Pilatus 6M detector at a sample-detector distance of 3 m and at a wavelength of *λ* = 0.124 nm. Data were collected in static mode (batch), as well as with an integrated SEC purification step, employing in-line size-exclusion chromatography. For this, an Agilent 1260 Infinity II Bio-inert LC, equipped with a GE Superdex 75 Increase 5/150 size exclusion column was used. 40 μL of sample were injected at a flow rate of 0.30 mL/min at 20°C. The column elute was directly streamed to the SAXS capillary cell. Approximately 1800 successive 0.5 s frames were collected. The data were normalized to the intensity of the transmitted beam and radially averaged, resulting in *I*(*s*) vs. *s* scattering profiles, where *s* = 4*π*sin*θ*/*λ* (momentum transfer), and 2*θ* is the scattering angle. The program CHROMIXS [22] was employed for visualization and reduction (background subtraction), both automatically and interactively, of the SEC-SAXS datasets. The PRIMUS [23] module of the ATSAS software package [24] was used for further processing. The radius of gyration (*R* _G_) was obtained by fitting the linear Guinier region of the data. Pair distribution function, *p*(*r*), with the corresponding maximum particle size parameter, *D* _max_), was determined using the program GNOM 7 [25]. Theoretical scattering profiles were computed from X-ray coordinates using Crysol [26]For the reconstruction of a theoretical molecular envelope, ab initio modeling, the program DAMMIF [27] was used, where scattering from the calculated envelopes was fitted against the experimental scattering and evaluated by the *χ* ^2^ values. For advance hybrid modeling, ensemble modeling was performed with EOM (Ensemble Optimization Method) 2.1 [28].

### 2.5 Circular dichroism

Far-UV CD spectrum of the human ^186-341^SF3B1 was recorded in the 180-260 nm wavelength range using a Jasco J-810 CD spectrometer. The protein sample was prepared at concentration of 6.8 µM in 5 mM ammonium acetate and 500 µM EDTA, pH 6.5. The corresponding buffer spectrum was recorded and subtracted to correct for background contributions. Measurements were performed at room temperature. Data points exceeding the instrument high-voltage threshold (600 V) were excluded. Baseline-corrected CD values were converted to molar ellipticity [Θ] according to standard procedures [29].

### 2.6 NMR Spectroscopy

The experiments were recorded on different spectrometers at 298 K: a Bruker Avance NEO spectrometer operating at 1200.64 MHz ^1^H, 301.93 MHz ^13^C and 121.67 MHz ^15^N frequencies (hereafter referred to as 1200 MHz spectrometer), equipped with a cryogenically cooled probe head optimized for ^13^C direct detection (CP2.1 TXO 1200SA N/C-H-D-05 Z XT); a Bruker Avance III spectrometer operating at 950.20 MHz ^1^H, 238.93 MHz ^13^C and 96.28 MHz ^15^N frequencies (hereafter referred to as 950 MHz spectrometer) equipped with a cryogenically cooled probe head (CP TCI 950S6 H-C/N-D-05 Z); a Bruker Avance NEO spectrometer operating at 899.50 MHz ^1^H, 226.20 MHz ^13^C and 91.16 MHz ^15^N frequencies (hereafter referred to as 900 MHz spectrometer) equipped with a cryogenically cooled probe head (CP TCI 900S6 H-C/N-D-05 Z); a Bruker Avance NEO spectrometer operating at 700.13 MHz ^1^H, 176.07 MHz ^13^C and 70.95 MHz ^15^N frequencies (referred to as 700 MHz spectrometer) equipped with a cryogenically cooled probe head optimized for ^13^C-direct detection (CP TXO 700S4 C/N-H-D-05 Z).

For ^186–341^SF3B1 protein ^15^N rates and heteronuclear ^1^H-^15^N NOEs [30] as well as CLEANEX [31] experiments were recorded on a sample of 320 µM ^15^N uniformly labelled ^186–341^SF3B1 in 50 mM sodium phosphate, 200 mM NaCl, 5 mM EDTA at pH 6.7. Experiments for the assignment and side chains characterization were recorded on a sample of 250 µM ^13^C^15^N uniformly labelled ^186–341^SF3B1 in the same buffer.

Backbone and side-chain assignments were obtained using a combination of ^1^H- and ^13^C-detected NMR experiments. The complete list of acquired experiments, and the most relevant acquisition parameters are reported in Table 1.

**Table 1.** Summary of NMR experiments acquired for sequential assignment of ^186-341^SF3B1 construct, together with the most relevant acquisition parameters.

| **Experiment** | **Field**<br>**(<sup>1</sup>H MHz)** | **Dimension of acquired data** |  |  | **Spectral width**<br>**(ppm)** |  |  | **NS** | **d1**<br>**(s)** |
|---|---|---|---|---|---|---|---|---|---|
|  |  | t<sub>1</sub> | t<sub>2</sub> | t<sub>3</sub> | F<sub>1</sub> | F<sub>2</sub> | F<sub>3</sub> |  |  |
| ***SF3B1*** |  |  |  |  |  |  |  |  |  |
| <sup>1</sup>H-<sup>15</sup>N HSQC [32] | 1200 | 2048 (<sup>1</sup>H) | 512 (<sup>15</sup>N) |  | 14 | 32 |  | 16 | 1.0 |
| CON [33] | 1200 | 2048 (<sup>13</sup>C) | 640 (<sup>15</sup>N) |  | 34.5 | 40 |  | 4 | 2.0 |
| HNCO [34] | 1200 | 2048 (<sup>1</sup>H) | 124 (<sup>15</sup>N) | 196 (<sup>13</sup>C) | 14 | 22 | 8 | 2 | 1.2 |
| HN(CA)CO [34] | 1200 | 2048 (<sup>1</sup>H) | 124 (<sup>15</sup>N) | 196 (<sup>13</sup>C) | 14 | 22 | 8 | 4 | 1.2 |
| CBCANH [35] | 1200 | 2048 (<sup>1</sup>H) | 128 (<sup>15</sup>N) | 236 (<sup>13</sup>C) | 14 | 22 | 60 | 4 | 1.2 |
| CBCA(CO)NH [35] | 1200 | 2048 (<sup>1</sup>H) | 128 (<sup>15</sup>N) | 236 (<sup>13</sup>C) | 14 | 22 | 60 | 4 | 1.2 |
| CACO [36] | 1200 | 2048 (<sup>13</sup>C) | 628 (<sup>13</sup>C) |  | 25 | 38 |  | 4 | 1.0 |
| CBCACO [37] | 1200 | 2048 (<sup>13</sup>C) | 826 (<sup>13</sup>C) |  | 25 | 75 |  |  | 1.2 |
| **(H)C**BCA(N)CO [38] | 700 | 1024 (<sup>13</sup>C) | 128 (<sup>13</sup>C) |  | 28.5 | 70 |  | 64 | 1.2 |
| (H)CBCACON [36] | 1200 | 2048 (<sup>13</sup>C) | 64 (<sup>15</sup>N) | 64 (<sup>13</sup>C) | 34.5 | 2 | 70 | 8 | 1.0 |
| (HCACON)N(CACO)NCO [39] | 1200 | 2048 (<sup>15</sup>N) | 96 (<sup>15</sup>N) | 128 (<sup>13</sup>C) | 25 | 40 | 40 | 8 | 1.3 |

All ^13^C-detected experiments were acquired using pulse sequences optimized for highly flexible protein regions. Homonuclear ^13^C-^13^C couplings were removed by applying an IPAP-based virtual decoupling scheme, in which in-phase and antiphase components were recorded and appropriately combined [40].

#### 2.6.1 ^15^N rates and heteronuclear NOEs

R_1_ relaxation rates were measured at 700 MHz using pseudo-3D ^1^H-^15^N correlation experiments, in which a series of 2D HN spectra were acquired with different relaxation delays stored along the third dimension. Data were collected with 8 scans (2024 × 512 points) and a recycle delay of 3 s, using relaxation delays of 22 ms, 60 ms, 150 ms, 250 ms, 400 ms, 600 ms, 800 ms, 1 s, 1.2 s, and 1.5 s. R_1_ values were obtained by fitting peak intensities to a single-exponential decay: *I* = *I*_0_ *e*^(–*R*^_1_*^t^*^)^.

R_2_ relaxation rates were measured on the same spectrometer using analogous pseudo-3D ^1^H-^15^N correlation experiments. Data were collected with 8 scans (2024 × 512 points) and a recycle delay of 3 s, using relaxation delays of 40.96 ms, 81.92 ms, 122.88 ms, 163.84 ms, 204.80 ms, 245.76 ms, 286.72 ms, 327.68 ms, 409.60 ms, and 491.52 ms. R_2_ values were obtained by fitting peak intensities to a single-exponential decay: *I* = *I*_0_ *e*^(–*R*^_2_*^t^*^)^.

^1^H-^15^N heteronuclear NOE [30] values were measured at 700 MHz from a 2D ^1^H-^15^N correlation experiment acquired with 64 scans (2024 × 448 points) and a recycled delay of 4.5 s. A reference spectrum was recorded without saturation and compared to a second spectrum acquired with ^1^H saturation. NOE values were calculated as the ratio of peak intensities in the spectra recorded with and without the saturation block.

#### 2.6.2 CLEANEX experiments

The CLEANEX experiments [31] were acquired at the 700 MHz spectrometer. The spectra were acquired with 32 scans (2048 × 200 points) and a relaxation delay of 1.5 s. The following delays were used: 5 ms, 10 ms, 20 ms, and 30 ms.

#### 2.6.3 Secondary structure propensity

Secondary structure propensities (SSP) were calculated using the ncSPC approach [41], based on backbone chemical shifts (H^N^, N, C^α^, C^β^, and C’).

#### 2.6.4 Temperature, pH, and ionic strength dependence experiments

The temperature coefficient of amide proton chemical shifts was measured by recording a series of 2D ^1^H-^15^N HSQC on the 900 MHz spectrometer at the following temperatures: 278 K, 283 K, 288 K, 293 K, 298 K, 303 K, and 308 K. All the experiments were acquired with 4 scans (2048 × 512 points). The pH dependence of amide proton chemical shifts was measured by recording a series of 2D ^1^H-^15^N HSQC on the 900 MHz spectrometer at the following pH values: 5.5, 6.0, 6.7, 7.3, and 8.0. All the experiments were acquired with 4 scans (2048 × 512 points). The response to ionic strength was assessed by recording a series of 2D ^1^H-^15^N HSQC on the 950 MHz spectrometer at the following NaCl concentrations: 0 mM, 50 mM, 100 mM, and 200 mM. All the experiments were acquired with 4 scans (2048 × 512 points) and a relaxation delay of 1.0 s.

#### 2.6.5 RNA titration

Protein-RNA interactions were investigated by NMR titration experiments using 150 μM of ^13^C,^15^N labelled ^186-341^SF3B1 sample in 50 mM sodium phosphate, 100 mM NaCl, 5.0 mM EDTA, at pH 6.3. The sample of protein was titrated with the RNA oligonucleotide up to 2.4 molar equivalents. Experiments were performed in 50 mM sodium phosphate, 100 mM NaCl, 5.0 mM EDTA, at pH 6.3. RNA-induced chemical shift perturbations were monitored using a combination of ^1^H- and ^13^C-detected NMR experiments, including ¹H-¹⁵N HSQC, CON, CACO and CBCACO spectra, acquired on a 700 MHz spectrometer at 298 K. While ^1^H-detected experiments provide high sensitivity, ^13^C-detected experiments offer improved spectral resolution and reduced signal overlap for IDPs, allowing reliable monitoring of residues affected by RNA interaction. In addition, experiments targeting selected side chains were employed to probe specific residue types.

#### 2.6.6 Data processing and analysis

All the spectra were acquired and processed using TopSpin 4.0 or above. The spectra were analyzed using TopSpin 4.0 and CARA (Computer Aided Resonance Assignment) [42]. The ^15^N R_1_ and R_2_ relaxation rates were determined by single-exponential fitting of cross-peak intensity decays as a function of the relaxation delay, using Bruker Dynamics Center 2.4.

### 2.7 Molecular dynamics (MD) simulations

A structural model of the wild-type ^186-341^SF3B1 fragment was obtained with AlphaFold3 [43] (Figure S1) From this, we built a model in which residues T211, T235, T313, T328 were phosphorylated. In both cases, the N- and C-termini were capped as NH₃⁺ and COO⁻, respectively. Protonation state of ionizable residues was determined with the Propka software [44]. We initially tested for the wild-type ^186-341^SF3B1 three different force fields: AMBER99SB-ILDN (A99-ILDN) [45] and CHARMM36m (C36m) [46]both with the TIP4P-D [47] water model, and the AMBER99SB-DISP (A99-DISP) [48] with the DISP water model. The phosphorylated ^186-341^SF3B1 was simulated only with the CHARMM36m force field, since this proved to better reproduce experimental data of the wild-type protein. The parameters of the phsophorylated threonines, holding a charge of −2e, were derived from reference [49]. Wild-type and phosphorylated ^186–341^SF3B1 were placed at the centre of dodecahedral simulation boxes, with a minimum distance of 3 nm between the protein and the box edges. Systems were solvated, neutralised with Na⁺ and Cl⁻ ions following the protocol of ref [48], and supplemented with NaCl to a final concentration of 200 mM to match the experimental ionic strength. Periodic boundary conditions were applied throughout. Bonds involving hydrogen atoms were constrained with the LINCS algorithm [50]. Short-range electrostatic and Lennard–Jones interactions used a 1.0 nm cutoff, while long-range electrostatics were treated with particle mesh Ewald (PME) using a grid spacing of 0.12 nm [51]. Each system was first energy-minimised using steepest descent. This was followed by a 100 ps NVT equilibration at 298 K, during which protein positional restraints of 1000 kJ mol⁻¹ nm⁻² were applied to allow relaxation of the solvent. Temperature was controlled using velocity-rescaling coupling with a coupling time constant of 0.1 ps [52]. The systems then underwent 500 ps of NPT equilibration with backbone restraints (1000 kJ mol⁻¹ nm⁻²), followed by 1 ns of unrestrained NPT equilibration. Pressure was maintained at 1 bar using the Parrinello–Rahman barostat with a coupling time constant of 2 ps [53]. Production simulations were run for 2 μs with a 2 fs timestep. For C36m and A99-DISP we performed 3 MD simulation replicas of 2 μs each, while for A99-ILDN only one replica of 2 μs was run, as this force field did not reproduce the experimental SAXS and NMR data well (vide infra).

MD simulation trajectories were analysed and compared with experimental observables, including SAXS profiles calculated using Pepsi-SAXS [54]. A99SB-DISP and C36m were selected for detailed comparison because they provided better agreement with the SAXS profiles. The SAXS-based R_g_ was computed using the Guinier approximation, applied to the low-q region of the experimental scattering profile. This approach provides a robust estimate of the overall size and compaction of disordered proteins in solution. The comparison between simulated and experimental R_g_ therefore offers a direct validation of the force field ability to reproduce ensemble-averaged chain dimensions. NMR chemical shifts were predicted using SPARTA+ [55].

We then evaluated how the ^186–341^SF3B1 fragment interacts with the RNA sequence that was shown to bind to ^186–341^SF3B1. Since both the RNA and ^186–341^SF3B1 fragments are disordered, we divided them into smaller fragments. To this end, the 20-nucleotide-long RNA was split in three 10-residue long sequences

RNA split 1: rCUACUACCCC (residues from 1 to 10)

RNA split 2: rACCCCCACCC (residues from 6 to 15)

RNA split 3: rCACCCUCACC (residues from 11 to 20)

Instead, for the ^186–341^SF3B1 protein we focused on the fragment surrounding W200 (i.e. fragment RKRRWDQTA, ^196-204^SF3B1), which NMR measurements demonstrated to interact with RNA. Additionally, we also considered, as a negative control, the PGRAK240GSET segment (^236-244^SF3B1), which instead did not interact with RNA in NMR experiments. To model SF3B1/RNA interactions we generated the structural ensembles of the adducts between each RNA split and ^196-204^SF3B1/^236-244^SF3B1 fragments using AlphaFold3 [43]. Briefly, we generated 5,000 structures per each RNA split (using 1000 seeds, each yielding 5 structures). Hydrogen atoms and terminal caps were added to each structure using the *tleap* module of AmberTools [56].

Interactions of ^196-204^SF3B1/^236-244^SF3B1 with the RNA splits were analysed with ProLIF [57]. The resulting interaction fingerprints were used as input features for dimensionality reduction and clustering analyses, enabling the identification of different initial bound states corresponding to distinct interaction patterns. Prior to clustering, we discarded structures exhibiting severe steric clashes (i.e. when any pair of non-bonded atoms were separated by less than 1.5 Å). We then performed clustering using HDBSCAN [58] with the Jaccard distance as the metric, given the binary nature of each feature. Features present in more than 95% or less than 5% of frames were excluded. For the ^196-204^SF3B1 fragment, we extracted two clusters for RNA splits 1 and 2 and three clusters for split 3. For each cluster we extracted two to three representative structures, giving a total of 7 initial conformations. For the ^236-244^SF3B1 fragment, clustering proved unfeasible, as the interaction landscape lacked sufficient diversity. Thus, in this case, we arbitrarily selected one representative structure for each split.

The selected structures were then used as starting configurations for MD simulations. Each system was solvated in a simulation box with a minimum distance of 1.8 nm between the solute and the box boundaries. Counter-ions (Na⁺ and Cl⁻) were added to neutralize the systems, together with additional monovalent ions to achieve an approximate ionic strength of 150 mM. Due to variations in the number of solvent molecules across systems and the use of a fixed number of ions, the actual salt concentration ranged between 130 and 150 mM.

In these MD simulations we employed Amber ff14SB force field [59] for the protein component, the χOL3 parameter set of parameters for RNA [60], in combination with the OPC water model [61]. since this force field combination is the most validated for describing protein/RNA systems. Monovalent ions were described using the Joung–Cheatham parameters [62].

## 3 Results

### 3.1 Small-angle X-ray Scattering Analysis

To gather structural information on the architecture of the human ^1–463^SF3B1 region, we carried out SAXS experiments on the whole N-terminal construct fused to a GST tag, since all attempts to remove the tag failed. To improve data quality, SEC-SAXS mode was employed, in which a size-exclusion chromatography column is directly connected to the SAXS measuring capillary, allowing separation of possible oligomeric species. Indeed, SEC-SAXS analysis revealed a single, symmetrical elution peak, indicating sample monodispersity (Figure S2A). The integrated SAXS intensity profiles (Figure S2B) and the corresponding background-subtracted scattering curves (Figure 2A) were of good quality and suitable for structural analysis. The GST-^1–463^SF3B1 fusion protein displayed a large Rg (6.6 ± 0.2 nm) and Dmax (20 ± 2 nm), consistent with the molecular dimension expected for the fusion construct. The broad pair-distance distribution functions P(r) and shifted Kratky profile (Figures 2B, C) suggest an extended particle, likely reflecting the contribution of the GST moiety and the longer polypeptide region (Table S2).

**Figure 2.**
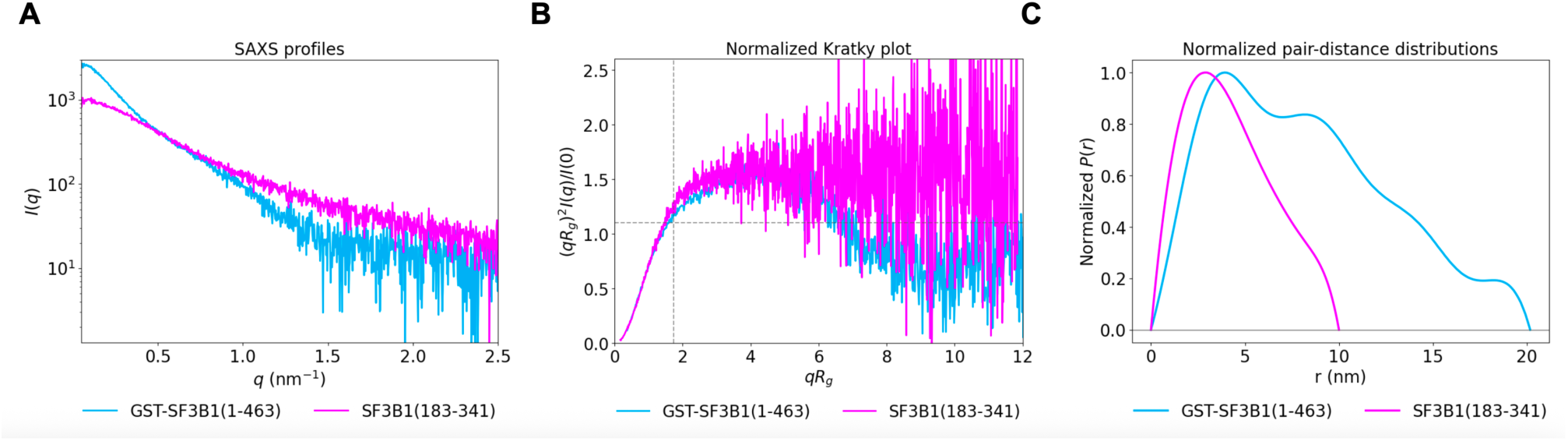
SAXS analysis of the ^186–341^SF3B1 peptide and GST-^1–463^SF3B1 fusion protein. (A) Background-subtracted SAXS scattering curves obtained by subtracting buffer frames from sample frames, displayed as log I(q) versus q. (B) Normalized Kratky plot. (C) Pair-distance distribution function, P(r), derived from the SAXS data.

We then analysed a ^186–341^SF3B1 fragment using SEC-MALS and SEC-SAXS. The UV elution profiles showed monodisperse peaks for all samples (Figure S2A), indicating the absence of major aggregation or heterogeneous species during data collection. The integrated SAXS intensity profiles (Figure S2B) and the corresponding background-subtracted scattering curves (Figure 2A) were of good quality and suitable for structural analysis. The ^186–341^SF3B1 constructs displayed a good scattering profile, and the Guinier analysis provided a radius of gyration (Rg) of 3.6 ± 0.1 nm, with maximum particle dimensions (Dmax) of 10.0 ± 1.0 nm. The normalized Kratky plots (Figure 2B) and the P(r), (Figure 2C), were consistent with those of an IDP (Table S2). Indeed, the theoretical Rg value expected for an IDP of the same length (155 amino acids) was estimated to be 3.5 nm, using Flory’s scaling law [63].

Far-UV CD analysis of the ^186–341^SF3B1 fragment was also performed. The resulting spectrum displayed a single pronounced minimum near 200 nm and lacked the characteristic signatures of α-helical (double minima at 208 and 222 nm) or β-sheet (minimum around 218 nm) secondary structures, indicating a predominantly random-coil conformation (Figure S3). These results support the classification of ^186–341^SF3B1 as intrinsically disordered region.

### 3.2 NMR assignment of the ^186-341^ SF3B1 construct: strategy and challenges

We next investigated the structural and dynamical properties of the ^186–341^SF3B1 by NMR spectroscopy. The backbone assignment was obtained by combining both ^1^H- and ^13^C-detected experiments (Figure S4). The ^1^H-detected experiments (including 2D ^1^H^15^N-HSQC and 3D HNCO, HNCACO, CBCANH, CBCACONH) provided a first set of correlations. However, the high abundance of sequence repeats, particularly the presence of 17 TP dipeptide repeats, led to severe chemical shift degeneracy, making an exclusively ^1^H-based assignment strategy inadequate. To overcome this limitation, additional ^13^C-detected experiments were acquired (including 2D CON, CACO, CBCACO and 3D CBCACON, (HCACO)N(CA)CON). Among these, the 3D (HCACO)N(CA)CON experiment, adapted from a higher dimensional (5D) experiment designed for IDPs [39], recorded at very high resolution represented a key step in the assignment strategy. By connecting two sequential CON cross peaks, it provided direct sequential connectivities. These connectivities were crucial for resolving the chemical shift degeneracy associated with the numerous TP repeats, enabling the linkage of ambiguous TP resonances and completing the backbone assignment (Figure 3).

**Figure 3.**
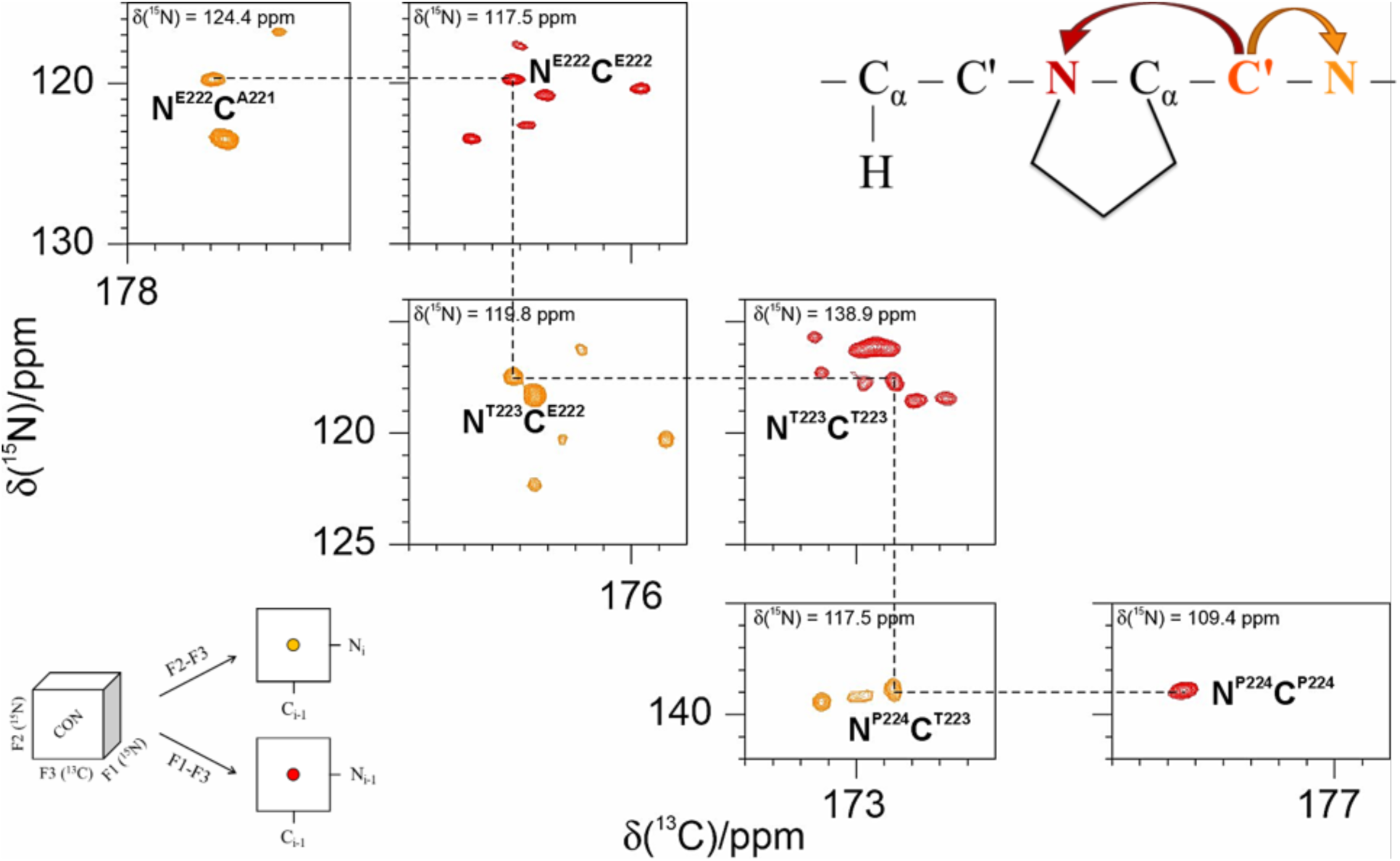
Representative spectra recorded for the wild-type ^186-341^SF3B1 construct. Representative planes extracted from the 3D (HCACO)N(CA)CON spectrum of ^186–341^SF3B1, illustrating sequential, residue-specific backbone assignment across proline-containing regions. The experiment correlates N_i_-C’_i_-N_i+1_, enabling sequential connectivity even in the presence of severe chemical shift degeneracy. Orange panels correspond to ^15^N_i+1_-^13^C_i_ correlations, and red panels correspond to ^15^N_i_-^13^C_i_ correlations; each panel is extracted at a defined chemical shift of ^15^N (indicated above it) in the 3D spectrum. Dashed lines highlight sequential correlations for the labelled residues (A^221^, E^222^, T^223^, P^224^), shown here as an example of how the experiment enables sequential assignment in regions representative of the many repeated motifs (TP, ETP, AETP) that occur throughout the protein.

By combining ^1^H- and ^13^C-detected datasets, an essentially complete backbone assignment was achieved (only the first two N-terminal residues of the construct remained unassigned). Backbone chemical shifts for H^N^, N, C’, C^α^, and C^β^ were obtained; in addition to the backbone, side-chain carbon resonances of N, D, E, and Q residues were assigned using 2D CACO and CBCACO spectra and those of the P with the 2D (H)CBCA(N)CO spectrum.

Protein dynamics and solvent accessibility was assessed through measurements of ^15^N longitudinal and transverse relaxation measurements, heteronuclear ^1^H-^15^N NOEs and CLEANEX experiments. The relaxation profiles showed values typical of an IDP, with no evidence of partially folded regions (Figure 4A). The protein displayed heterogeneous distribution of exchange with the solvent properties along the sequence, with an overall trend of increasing signal intensities while increasing the mixing times, consistently with widespread solvent exposure (Figure 4B). A subset of residues exhibited comparatively lower exchange, possibly due to more bulky side chains (such as tryptophan) as well as transient local interactions.

**Figure 4.**
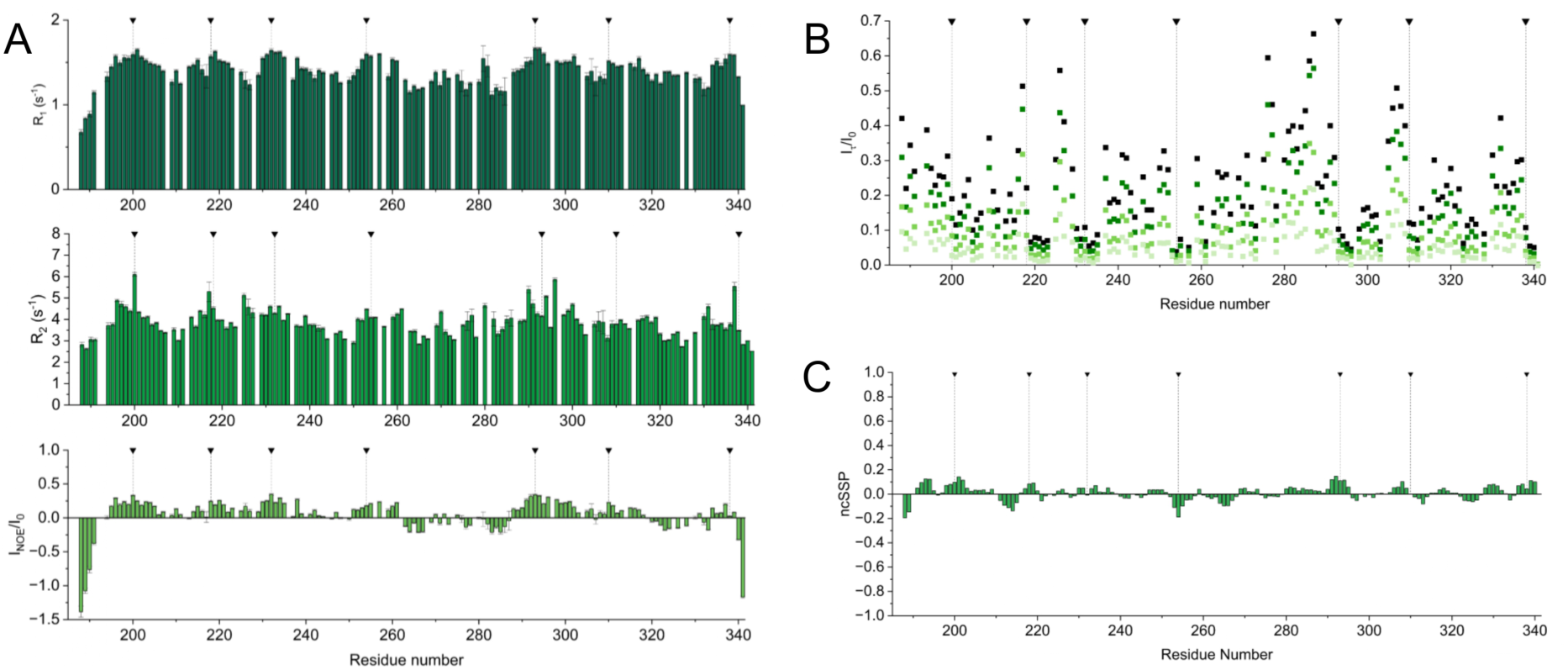
NMR Characterization of the wild-type ^186-341^ SF3B1 construct. (A) ^15^N relaxation rates, together with heteronuclear ^1^H-^15^N NOE values, are shown for the ^186–341^SF3B1 protein. (B) CLEANEX intensity profiles at different mixing times, normalized to the reference spectrum intensity, plotted as a function of residue number. Data are shown for mixing times of 5, 10, 20, and 30 ms, with colours ranging from light (shorter mixing time) to dark (longer mixing time). (C) Secondary structure propensity (SSP) profiles, calculated from backbone chemical shifts using ncSPC [41], shown as a function of residue number for the ^186–341^SF3B1 construct. Black triangles with dashed vertical lines indicate the positions of tryptophan residues.

Residue-specific secondary structure propensities (SSP) calculated from the NMR chemical shifts of the backbone nuclei (Figure 4C) confirmed the high disorder of the protein, with minimal helical or β-sheet character. Interestingly, slightly higher SSP values are observed for a small subset of polypeptide segments, including R^199^-W^200^, R^231^-W^232^, R^292^-W^293^, and R^337^-W^338^, suggesting a locally increased propensity for α-helical conformations. These sites coincide with aromatic tryptophan residues preceded by positively charged side chain residues, corresponding to regions of potential functional relevance.

The NMR characterization was completed determining the protein behaviour upon changing temperature, pH, and ionic strength (Figure S5). Most residues generally displayed linear temperature-dependent perturbations, with comparable chemical shift differences. An exception occurred at positions immediately following tryptophan residues. In six of the seven W sites, the residue immediately following the W side chain exhibited markedly attenuated chemical shift changes. Only one tryptophan site (W^254^, within the IWDP motif) showed a distinct behaviour, with higher chemical shift differences. Variation of pH from 5.5 to 8.0 revealed limited changes, with most NMR signals chemical shifts remaining nearly invariant. Only a subset of residues showed appreciable perturbations, particularly several resonances of histidine and serine residues progressively broadened or disappeared at higher pH, suggesting an enhancement of the exchange with solvent typical for these residues. The response to ionic strength was assessed by increasing NaCl concentration from 0 to 200 mM in buffer solution composed of 50 mM sodium phosphate and 5 mM EDTA at pH 6.7. Most residues are unaffected by the variation, while a minority exhibited minimal salt-dependent chemical shift changes, indicating local electrostatic effects.

Thus, NMR data demonstrate that ^186-341^SF3B1 behaves predominantly as a highly flexible, disordered protein segment lacking a stable folded structure, although a few short segments surrounding tryptophan residues exhibit a slight propensity to adopt transient local structure, which may contribute to its biological function.

### 3.3 Molecular Dynamics simulations

Since the reliability of MD simulations in capturing the structural dynamics of IDPs remains challenging, we initially simulated the wild-type ^186–341^SF3B1 fragment using different force fields and we monitored the agreement of the resulting conformational ensemble with the experimental SAXS and NMR data detailed above. To this end, we used the Amber99SB-ILDN (A99-ILDN) [45], CHARMM36m (C36m) [46] both with the TIP4P-D water model [47], and Amber99SB-DISPforce field with the DISP water model (A99-DISP) [48], which are most commonly employed to study IDPs. To monitor the agreement between the MD-derived and experimental SAXS curves, we compared the experimental scattering intensity profiles with the SAXS curves computed from the MD simulations. We observed that the C36m force field reproduced the experimental SAXS profile remarkably well, yielding the lowest discrepancy among the three force fields (χ^2^=1.04). The computed curve closely follows the experimental data, indicating that the conformational ensemble generated by using C36m force field is consistent with SAXS measurements (Figure S6).

A99-DISP also provides a good description of the SAXS data, although with a somewhat larger deviation (χ^2^=1.64). The overall shape of the scattering profile is reproduced, but subtle differences suggest an ensemble that is slightly more compact than that observed experimentally.

Instead, the A99-ILDN shows the poorest agreement (χ^2^=6.23). The calculated SAXS curve deviates substantially from the experimental profile, indicating that the simulated ensemble does not capture the overall dimensions and shape of the protein in solution.

We also computed distribution of the Radius of Gyration (Rg) on MD simulation trajectories (Figure S6), confirming that C36m and A99-DISP force fields provide a better description of the experimental scattering profiles than A99-ILDN, with C36m better aligning to the experimental values than A99-DISP.

An independent assessment of force field reliability was also achieved by comparing experimentally derived NMR chemical shifts with the (Figure S7) chemical shifts computed from the MD simulations. The resulting global root mean square error (RMSE), which quantifies the overall difference between the predicted values and experimentally observed chemical shifts (excluding amide-nitrogen shifts), was 0.437 ppm for C36m, 0.476 ppm for A99-DISP, and 0.485 ppm for A99-ILDN. These results demonstrated that C36m offers the most accurate representation of the ^186–341^SF3B1 conformational ensemble, with A99-DISP providing the next best agreement with the experimental data (Figure S7).

As both C36m and A99-DISP force field provided a more accurate description of the experimental SAXS and NMR observables, we performed three independent MD simulation replicas with these two force fields and we computed the correlation between experimental chemical shifts for different nuclei with those back-calculated from the combined trajectories (Figure S8). This analysis consistently showed that the conformational ensemble generated with C36m is in closer agreement with the NMR measurements. The superior performance of C36m in reproducing both SAXS and NMR observables motivated its selection for subsequent MD simulations of the phosphorylated ^186–341^SF3B1 variant. Thus, 3 MD simulation replicas of ^185–341^SF3B1 fragment, phosphorylated at pT211, pT235, pT313, and pT328, were done using the C36m force field.

Interestingly, the R_g_ and the distribution of the end-to-end distances (R_ee_) computed on the collective MD trajectories of the phosphorylated variant revealed a more compact conformational ensemble than the wild-type counterpart (Figure 5). To compare the level of compactness of the ^185–341^SF3B1 fragment in the wild-type and phosphorylated states, we inspected their solvation properties by computing the per-residue solvent-accessible surface area (SASA) (Figure S9). Consistently with the more compact conformational ensemble inferred from the R_ee_ and R_g_ distributions, the computed SASA revealed that phosphorylated variant had lower solvent exposure than the wild-type one. This is due to the presence of a larger number of contacts in the ^186–341^SF3B1 phosphorylated variant (Figure 5). In the wild-type ensemble, hydrophobic interactions were identified (Figure S10) with the most persistent contacts involving the tryptophan residues W218, W200, W218, W232, W254, W310, and W338, which interact either with one another or with the neighbouring isoleucine and leucine residues. Upon phosphorylation, this interaction network is substantially remodelled (Figure S10) leading to a small decrease in hydrophobic interactions and a marked increase in the number of intramolecular hydrogen bonds and salt bridges involving the phosphorylated pT211, pT235, pT313, and pT328 residues. These interactions contribute to the observed compaction of the phosphorylated conformational ensemble.

**Figure 5.**
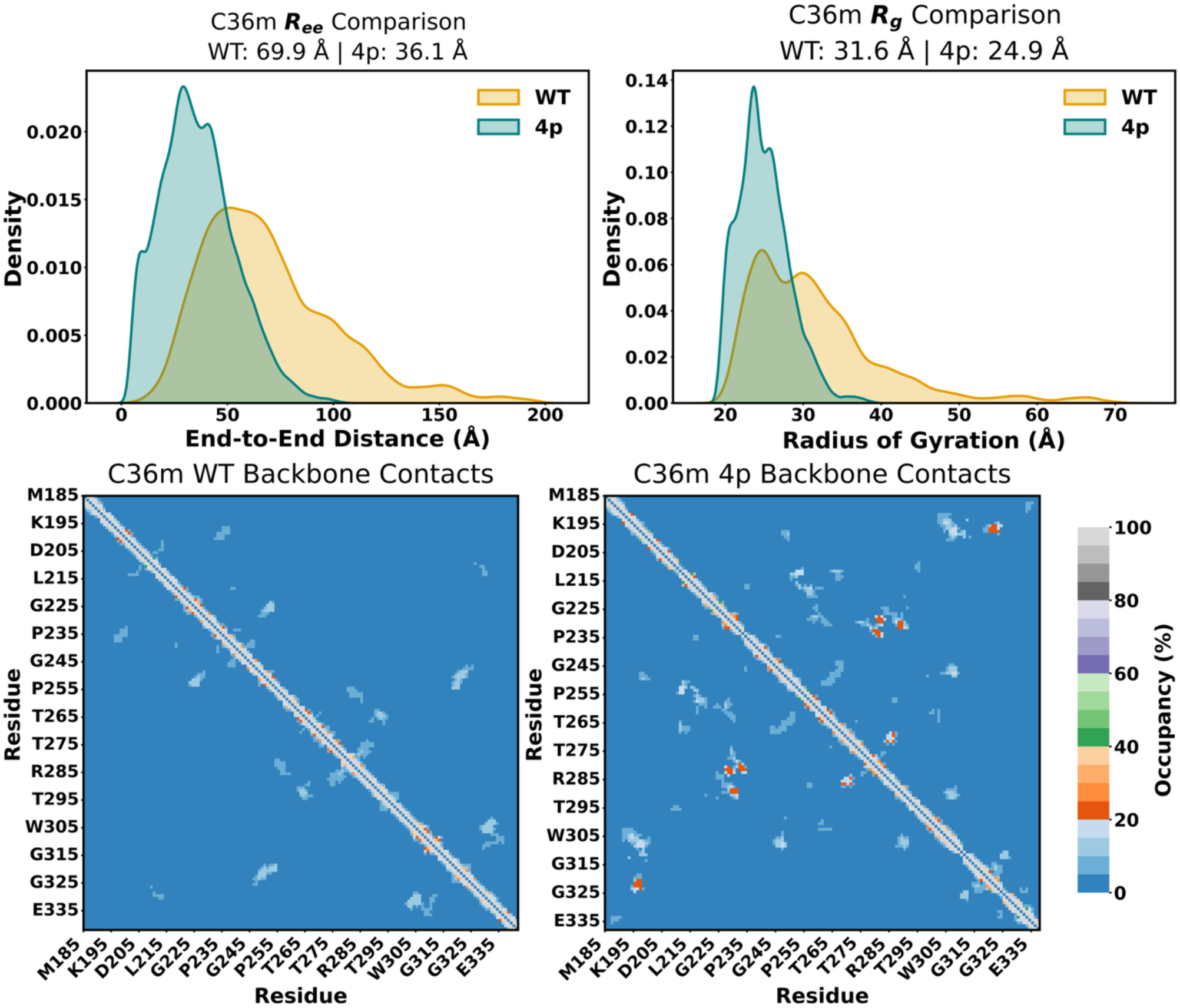
Structural Ensemble and Intramolecular Interactions of the wild-type and phosphorylated ^86-341^SF3B1. Top panel. Distribution of end-to-end distance (R_ee_, Å) and of the radius of gyration (R_g_, Å) as obtained from molecular dynamics simulations, performed with the CHARMM-C36m (C36m) force field. Data are reported for the wild-type (gold) and phosphorylated 4p (green) ^186–341^SF3B1 fragment. Bottom panel. Intramolecular contact probability of the ^186–341^SF3B1 IDR. Backbone contact maps (6 Å cutoff) are shown for wild-type and phosphorylated states across the MD simulations performed with the C36m force fields.

### 3.4 Bioinformatic analysis of SF3B1 IDR primary sequence

In contrast to the typical sequence of IDPs/IDRs, primary sequence of ^186–341^SF3B1 contains an unusually high abundance of tryptophan residues [64,65]. Specifically, nine W residues are present within the entire SF3B1-NTD, with seven of them being clustered within the ^186–341^SF3B1 fragment. These seven W residues occur in similar local sequence contexts, characterized by basic residues upstream (K or R) and acidic or polar (D, E, T) residues downstream (Table 2). Comparison of the ±4 residues surrounding each tryptophan reveals a recurring sequence pattern. These residues indeed are not distributed randomly, but organized into repeated local motifs, which conform to the well characterized U2AF Binding Motifs (ULM) for proteins containing U2AF Homology Motifs (UHM) [14,15].

**Table 2.** Sequence context of the seven W residues within the ^186–341^SF3B. fragment. For each W, the number of the residue and the four amino acids preceding and following it are reported.

|  |  |  |  |  |  |  |  |  |
|---|---|---|---|---|---|---|---|---|
| **R** | **K** | **R** | **R** | **W<sup>200</sup>** | **D** | **Q** | **T** | **A** |
| **K** | **L** | **S** | **S** | **W<sup>218</sup>** | **D** | **Q** | **A** | **E** |
| **P** | **S** | **L** | **R** | **W<sup>232</sup>** | **D** | **E** | **T** | **P** |
| **G** | **S** | **K** | **I** | **W<sup>254</sup>** | **D** | **P** | **T** | **P** |
| **R** | **K** | **N** | **R** | **W<sup>293</sup>** | **D** | **E** | **T** | **P** |
| **H** | **G** | **S** | **G** | **W<sup>310</sup>** | **A** | **E** | **T** | **P** |
| **R** | **K** | **S** | **R** | **W<sup>338</sup>** | **D** | **E** | **T** | **P** |

Thus, to assess the evolutionary conservation of these features, sequence alignments of the whole SF3B1 protein were done using orthologous proteins from different species (including *human*, *mouse*, *Schizosaccharomyces pombe*, *Xenopus laevis*, and *Saccharomyces cerevisiae)*. This comparative analysis revealed that while the folded CTD of SF3B1 is conserved across species, the disordered NTD exhibits a higher degree of variability. This pattern is consistent with the general behaviour of IDRs, which typically evolve more rapidly than structured globular domains [66]. Phylogenetic analyses of the NTD further support this domain-specific evolutionary pattern (Figure S11A). Despite the pronounced sequence divergence of the SF3B1-NTD across species, comparison of the two most distantly related orthologues revealed the persistence of several short-conserved sequence motifs (Figure S11B). This observation suggests that, although the NTD is disordered and rapidly evolving, selective pressure is maintained on discrete functional elements. Notably, these conserved ULM motifs are known to mediate interactions with the UHM-containing proteins, such as SF1 and U2AF2 [14] and are thus expected to act as interaction hotspots (Figure S11). Consistently with these observation, binding studies showed that, with the exception of the ULM motifs containing W254 and W310, which do not bind U2AF2 [14], all remaining ULMs interact with the U2AF2 UHM domain, albeit with different affinities (K_d_= 6.0 ± 4.7, 9.5 ± 9.2, 15.8 ± 8.7, 21.8 ± 13.5, 36.5 ± 22.6 mM for W338 > W293 > W200 > W218 > W232, respectively [14].

### 3.5 Biochemical characterization of SF3B1-NTD-RNA interactions

Since the spliceosome proteins are enriched in IDRs [2,3], which can mediate interactions with RNA strands [67], we also investigated the ability of the ^1–463^SF3B1 and ^186–341^SF3B1 fragments to interact with different nucleic acid models. We first assessed the binding properties of the GST-tagged ^1–463^SF3B1 fragment with different types of DNA and RNA substrates, including ssRNA, ssDNA, DNA fork, DNA bubble, D-loops, and R-loop structures, using Electrophoretic Mobility Shift Assays (EMSA) (Figures 6A, B). The ^1–463^SF3B1 fragment displayed higher affinity for R-loops, RNA fork and ssRNA substrates, followed by D-loops (Figure 6C). GST alone did not bind ssRNA under the same experimental conditions (Figure S12) confirming that the observed interactions were mediated by the SF3B1 fragment. In contrast, the ^1-463^SF3B1 fragment showed substantially lower affinity for the other DNA substrates tested (i.e., DNA fork, DNA bubble, and ssDNA) (Figure 6C).

**Figure 6.**
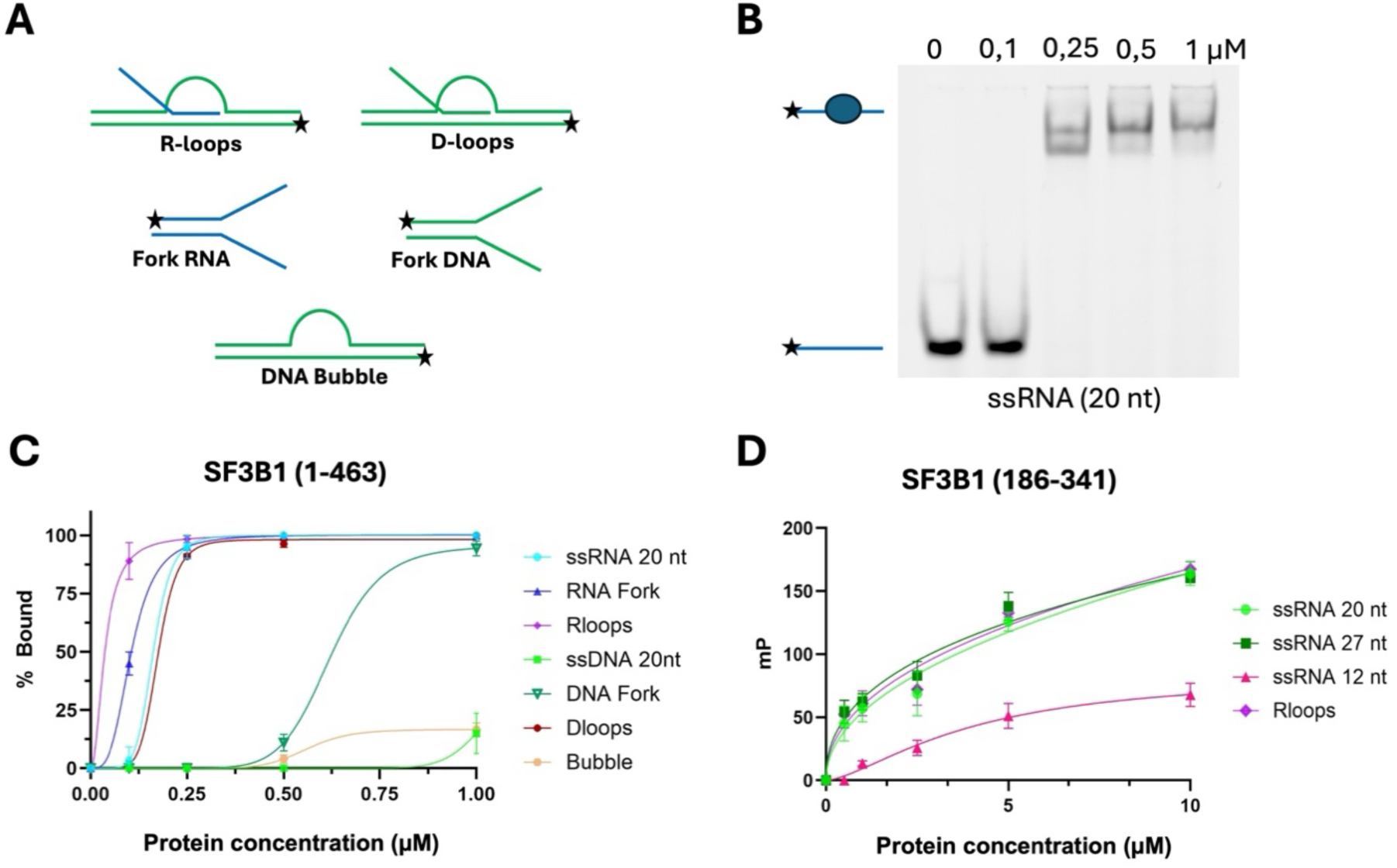
Biochemical analysis of ^1-463^SF3B1 and ^186-341^SF3B1 fragment. (A) Schematic diagram of the substrates used. A star denotes the position of the fluorescent 6-FAM moiety. (B) Example of a gel shift assays for the ^1–463^ SF3B1 N-terminus fragment using a single strand (ss)RNA. The assays were carried out at increasing concentrations of protein (0–1 μM); each experiment was done in triplicates. (C) Comparison of ^1–463^SF3B1 binding affinity towards a variety of DNA and RNA substrates. The protein shows a preference for R-loops, closely followed by RNA fork, ssRNA and D-loops. (D) Fluorescence anisotropy measurements for the binding of ^186–341^SF3B1 fragment to ssRNA with different lengths and R-loops. The protein has a significant lower affinity when compared to the whole N-terminal domain. A binding analysis based on the one-site specific binding with Hill slope model.

Based on these results, we next investigated the interaction of the ^186–341^SF3B1 fragment with the same nucleic acids models. Since EMSA experiments did not yield reproducible binding signals for this fragment, potentially reflecting the highly disordered nature of the ^186–341^SF3B1 region and the resulting instability or altered electrophoretic migration of protein–nucleic acid complexes, binding was assessed using fluorescence anisotropy measurements [68]. These experiments indicated that ^186–341^SF3B1 binds both ssRNA substrates and R-loops, although with lower affinity compared to the full ^1–463^SF3B1-NTD (Figure 6D).

### 3.6 NMR experiments elucidate ^186–341^SF3B1 binding to ssRNA

To validate and complement these insights, we monitored the interaction of ^186–341^SF3B1 with a 20 nucleotide-long ssRNA sequence by recording ¹H-¹⁵N HSQC and CON spectra of ^186–341^SF3B1 in the presence of increasing quantities of RNA. Chemical shift perturbations (CSP), together with normalized intensity differences and intensity ratio, identified a subset of residues affected by RNA binding (Figure 7). NMR titration data indicated that RNA binding primarily affected residues 189– 202 of the ^186–341^SF3B1 construct. The consistent response of this region in both CSPs and signal intensities suggests a key role in RNA binding. The increase in ^1^H–^15^N HSQC intensity is consistent with reduced solvent exchange and increased protection of amide protons upon RNA binding. Conversely, the decrease in CON intensity is consistent with an increase in the local effective rotational correlation time (τ_r_), resulting in enhanced transverse relaxation and signal broadening. Together, these data support a direct interaction of residues 189–202 (^189^ASQPPSKRKRR**W**DQ^202^) with RNA substrates.

**Figure 7.**
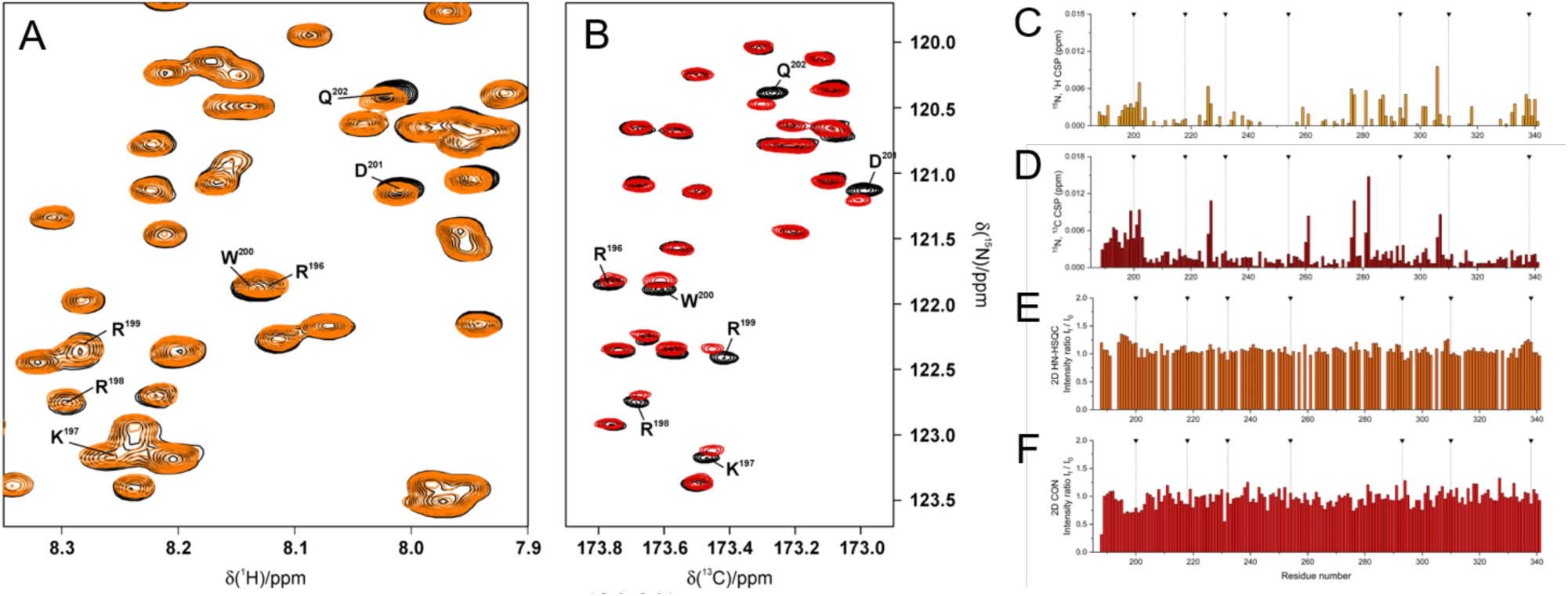
NMR characterization of ^186–341^SF3B1-RNA interaction. (A) 2D ¹H-¹⁵N HSQC and (B) 2D CON spectra of the ^186–341^SF3B1 protein with RNA (150 μM free protein and + 2.4 equivalents of RNA). The reference spectra are shown in black, while the ones acquired in presence of RNA are shown in orange and red for 2D ¹H-¹⁵N HSQC and 2D CON spectra respectively. Residue-specific chemical shift perturbations (CSP) derived from 2D ¹H-¹⁵N HSQC (C) and 2D CON (D) spectra, and residue-specific intensity ratio (RNA-bound/free protein) obtained from 2D ¹H-¹⁵N HSQC (E) and 2D CON (F) spectra. Data highlight residues involved in RNA binding. Black triangles with dashed vertical lines indicate the positions of tryptophan residues.

From a sequence perspective, this RNA-interacting region of ^186–341^SF3B1, besides exhibiting a tryptophan residue, is enriched in positively charged amino acids, particularly in lysine and arginine residues, which can establish favourable electrostatic interactions with the negatively charged RNA phosphate backbone. This support an RNA recognition mechanism by ^186–341^SF3B1 involving both electrostatic and aromatic contacts, a common feature of protein-RNA interfaces [69].

### 3.7 Predicting ^186–341^SF3B1 RNA interactions via MD simulations

We performed MD simulations to characterize the interactions between ^186–341^SF3B1 and the RNA sequence used in the NMR study. To this end, we adopted a fragment-based strategy, which allows exploring the interactions between the two partners, while retaining the essential molecular features underlying their recognition and binding. Namely, the 20-nt RNA was divided into three overlapping segments (see Methods), while the ^186–341^SF3B1 was represented by a 9-residue long RNA-binding motif centred on W200 (^196^RKRRWDQTA^204^, ^196-204^SF3B1), since NMR experiments revealed that this segment best interacts with ssRNA. Additionally, as a negative control, we also considered a non-interacting 9-residue long peptide centred on K240 (^236^PGRAKGSET^244^, ^236-244^SF3B1) since this part of ^186–341^SF3B1 was instead not observed to interact with RNA in NMR experiments.

Briefly, structural ensembles of ^196-204^SF3B1/RNA adducts were generated with AlphaFold3 and analysed via interaction fingerprints to capture the diversity of ^196-204^SF3B1/RNA contacts. Clustering of interaction patterns revealed multiple distinct binding modes for the W200-containing SF3B1 fragment, which resulted into seven representative initial conformations considering all RNA splits (Figures S13-S14). In contrast, the ^236-244^SF3B1 region displayed a limited interaction landscape. Consequently, the conformational ensemble could not be readily partitioned into clusters, and a representative structure was therefore selected arbitrarily for each split.

Starting from these representative structures, 1 μs-long MD simulations in explicit water were performed to relax the adducts under near-physiological conditions. Notably, the contacts between ^196-204^SF3B1 and ^236-244^SF3B1 and each RNA split initially present in the adduct models rapidly changed and, in some cases, the IDR/RNA, adduct dissociated within the first 200 ns of MD simulations (Figure S15-S16).

We then characterised the interaction patterns between RNA and ^196-204^SF3B1 to pinpoint the molecular basis of this interaction. This analysis revealed that the interaction between the W200-centered fragment and the RNA strands is primarily mediated by three R residues, which establish hydrogen bonding, cation–π, and salt bridges interactions with the RNA phosphate backbone. W200 also contributed significantly to binding by establishing π–π-stacking interactions with the RNA bases. However, from an energetic standpoint, cation–π interactions and salt bridges are generally stronger than π–π stacking interactions, suggesting a dominant role for R-mediated contacts in stabilising the RNA/IDR complex (Figure S16).

To gather an overall picture of the interaction mode between the W200-centered SF3B1 segment and the RNA splits, we computed residue-wise average interaction frequencies across all simulations (Figure 8). This analysis confirmed that binding of ^196-204^SF3B1 to RNA is primarily driven by the three R residues preceding W200, with also W200 contributing to RNA binding.

**Figure 8.**
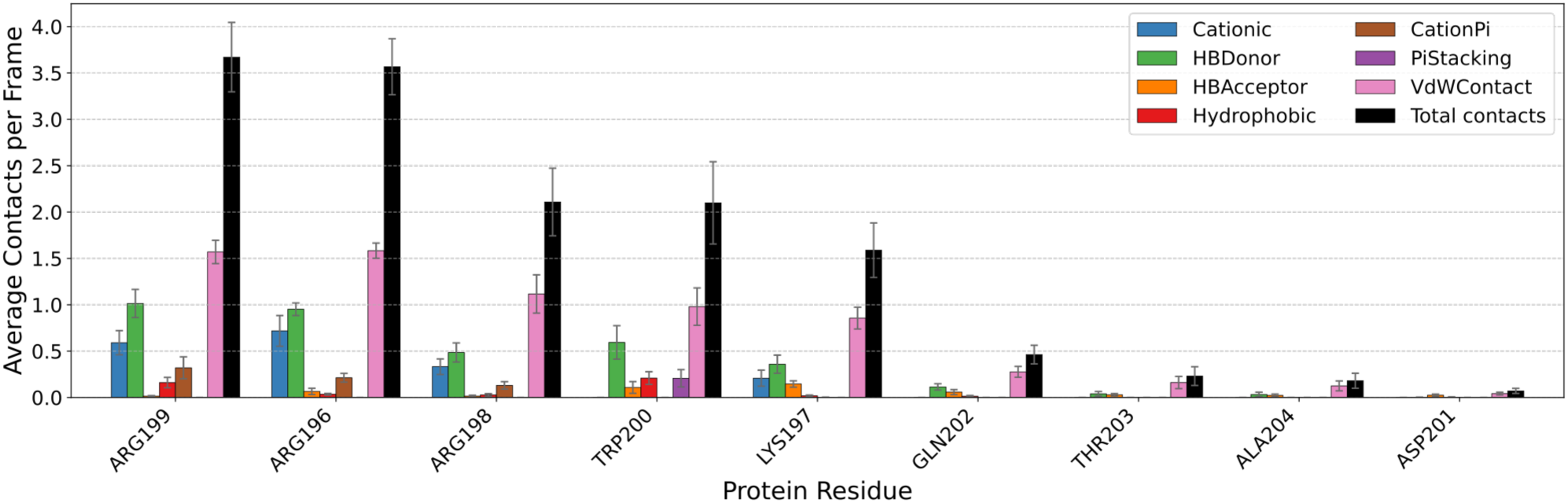
Nature and frequency of interactions between ^196-204^SF3B1 and RNA. Average interactions per residue of the RKRRW200DQTA fragment (^196-204^SF3B1) computed over all simulations across all RNA splits and clusters. Error bars represent the standard deviation calculated over all MD simulations, divided by √(N−1) (with N = 7). The first 100 ns of each trajectory were excluded from the analysis.

Although the interaction patterns between ^196-204^SF3B1 and RNA varied considerably among replicas, R residues consistently played a dominant role in binding. Their recurrent involvement in hydrogen bonds and salt bridges indicates that electrostatic interactions are the main determinants of ^196-204^SF3B1/RNA recognition. Additionally, ^196-204^SF3B1 binding to the RNA segments is highly non-specific (Figures S17-S18). Indeed, analysis of the MD trajectories did not reveal a clear preference for specific nucleobases or RNA sequence motifs (Figure S19). Rather, sequence selectivity appears to be governed by broader physicochemical features, including purine versus pyrimidine identity, intrinsic base-stacking propensities, and the availability of nucleobases capable of engaging in π-stacking interactions, either with neighbouring bases or with aromatic and cationic residues such as W and R. Despite the largely non-specific binding observed, in MD simulations performed with the negative control (the K240-containing peptide ^236-244^SF3B1) the contacts between the RNA and ^236-244^SF3B1 present in the initial adduct models were rapidly lost (Figures S20), consistently with the lack of interactions observed in NMR experiments.

Overall, this integrative approach highlights the heterogeneous and transient nature of SF3B1–RNA interactions, with the W200-centered ULM motif displaying diverse binding modes. These results support a model in which RNA recognition by SF3B1 IDRs is mediated by sequence-dependent interaction hotspots embedded within a highly dynamic conformational landscape (Scheme 1).

## Discussion

Through an integrated biophysical, biochemical, spectroscopic, and computational framework, we have elucidated the structural, and dynamical properties of a regulatory IDR fragment of the SF3B1 protein, a core component of the SF3b splicing factor. First SAXS experiments revealed that the whole SF3B1-NTD is intrinsically disordered, consistently with previous studies [14,15] and that it exhibits a highly dynamic conformational ensemble in solution.

Next, specifically focusing on the ^186−341^SF3B1 region, which contains four regulatory phosphorylation hotspots, we confirmed its intrinsically disordered nature via SAXS and NMR experiments. These experimental data were subsequently employed to benchmark MD force fields, revealing that C36m force field provides the best overall agreement with the experimental observables. Thus, this force field was selected to perform subsequent MD simulations aimed at predicting the structural consequences of phosphorylation on the ^186−341^SF3B1 conformational behaviour. MD simulations revealed that phosphorylation induces a significant compaction of the ^186−341^SF3B1 ensemble, driven by the formation of intramolecular salt bridges and hydrogen bonding interactions. Consistently, phosphorylation has been shown to remodel the conformational ensembles of IDR, by promoting substantial chain compaction [70,71].

In its unphosphorylated state, the SF3B1-NTD has been reported to interact with multiple UHM-containing splicing factors, including SF1, U2AF65, PUF60, TAT-SF1, RBM39, and SPF45 (RBM17) [14,15,72] [73]. Our findings provide a mechanistic framework for the phosphorylation-dependent regulation of the SF3B1 IDR. Specifically, they suggest that phosphorylation may either prevent the association of the SF3B1-NTD with U2AF2 or promote dissociation of preformed SF3B1-U2AF2 complexes. This model is supported by recent crystallographic studies of SF3B1 peptides encompassing W293 and W338-centered ULMs. These strudies revealed that phosphorylation of the SF3B1-NTD introduces negatively charged phosphoryl groups that are electrostatically incompatible with an acidic α-helix of U2AF2, thereby creating repulsive interactions that destabilize the SF3B1-U2AF2 complex and facilitate its dissociation [16]. As such, by reshaping its conformational landscape and interaction profile, phosphorylation is most likely poised to control the network of molecular contacts established by SF3B1 within the protein components of the spliceosome. Since phosphorylation is highly dynamic and reversible, such modifications of the SF3B1-NTD likely act as molecular switches that govern the spatiotemporal recruitment/release of splicing factors during spliceosome assembly, activation, and catalysis.

The several evolutionarily conserved W-centred motifs, clustering within the ^186-341^SF3B1 segment are key to mediate transient molecular interactions with several splicing factors. However, here we extend this model by demonstrating that the SF3B1-NTD is also capable of binding RNA, with a preference for R-loops. Combining RNA-binding experiments, NMR spectroscopy, and MD simulations, we show that the W200-centred region interacts with RNA through a predominantly nonspecific, yet sequence-dependent mechanism driven by a network of R- and W-mediated contacts. The resulting protein-RNA complexes remain highly dynamic, supporting a model in which the SF3B1 IDR enables adaptable and reversible interactions, while preserving selective molecular recognition. Interestingly, the W200-centered ULM exhibits weaker affinity for U2AF2 as compared to the W338 and W293-ceneted ones [14]. The W200-centerd motif exhibits a unique charge distribution among the W-containing ULMs, being flanked by five basic residues (KRKRR) and followed by only a single acidic residue (D). The pronounced local positive charge is expected to favour interactions with the negatively charged RNA backbone. Together, these observations suggest that the SF3B1-NTD may play a dual role during 3′ splice-site selection, mediating both protein-protein and protein-RNA interactions. Hence, the intrinsically disordered nature of the SF3B1-NTD is consistent with a scaffolding function that enables the formation of multiple, transient interaction networks. These interactions can be dynamically modulated by phosphorylation, which reshapes the conformational landscape of the SF3B1-NTD and thereby alters its binding preferences and regulatory functions [72].

**Scheme 1.**
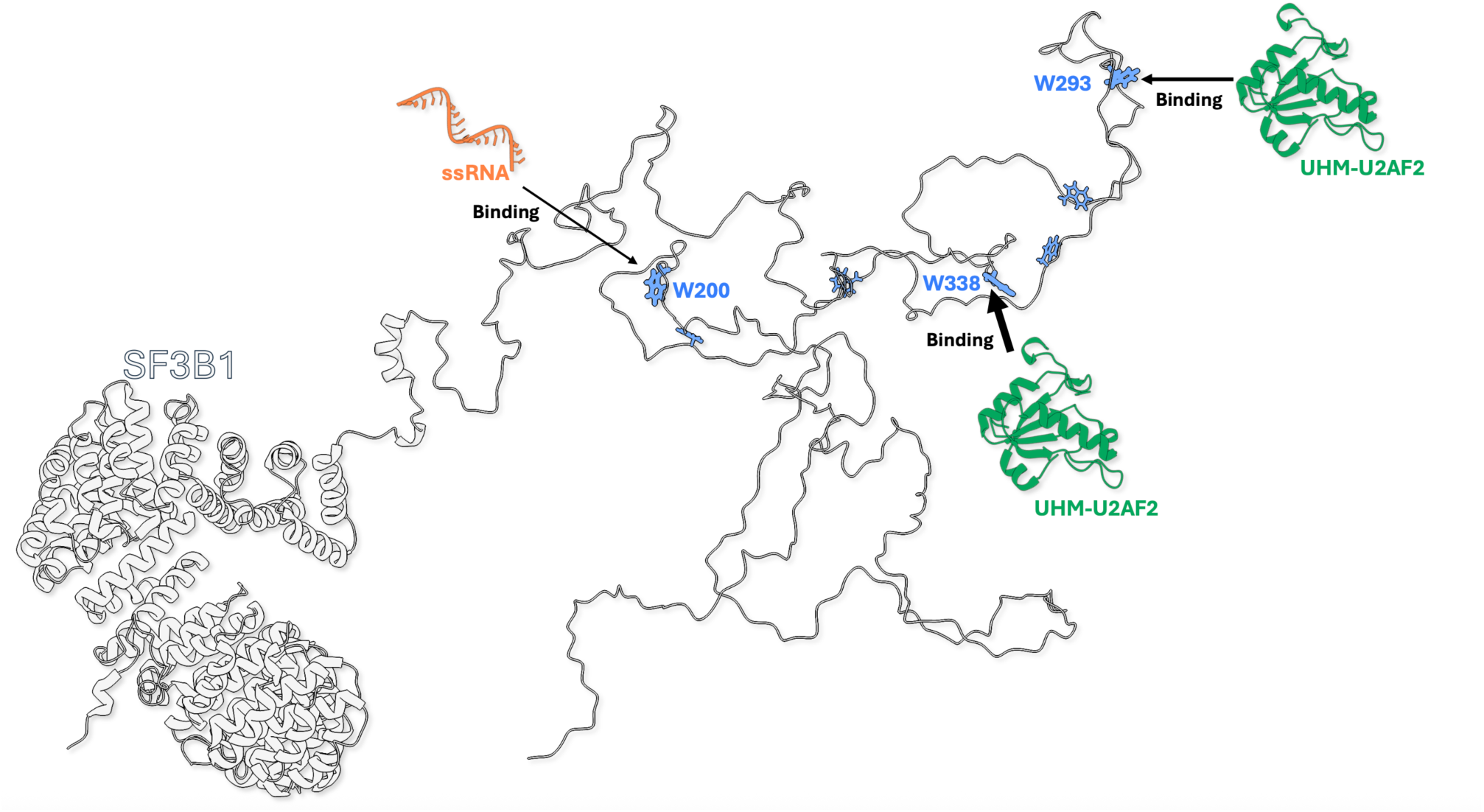
Scheme of the possible interactions of SF3B1-NTD with RNA strands and the UHM containing protein U2AF2.

Notably, SF3B1 mutations have been reported to induce R-loop accumulation in chronic lymphocytic leukemia [74]. Thus, it is tempting to suggest that the preferential interactions of SF3B1-NTD with these RNA sequences may take part in this process.

Altogether, our findings provide an example of how intrinsic disorder within SF3B1 encodes the regulatory and interaction capabilities required for spliceosome function. More broadly, the integrative experimental–computational strategy presented here provides a robust framework for investigating spliceosome IDP/IDRs and contributes to a deeper understanding of their roles in RNA-rich cellular environments.

## Conclusions

Our integrative approach combined SAXS, NMR, RNA-binding experiments and MD simulations establishes the SF3B1 NTD as a highly dynamic intrinsically disordered regulatory region whose structural and functional properties are finely modulated by phosphorylation. We show that phosphorylation induces significant compaction of the ^186−341^SF3B1 segment through the formation of intramolecular electrostatic interactions, providing a mechanistic basis for the regulation of its interactions with spliceosome proteins. We also demonstrate that the ^186−341^SF3B1 can directly bind RNA at its conserved W200-centered motifs, by establishing electrostatic and π–π-stacking interactions. This reveals a previously underappreciated role of SF3B1-NTD in RNA recognition. These findings support a model in which the SF3B1 IDR acts as a multifunctional interaction platform that dynamically balances protein-protein and protein-RNA interactions during spliceosome assembly and function. More broadly, our work highlights how intrinsic disorder and reversible phosphorylation cooperate to regulate the interaction landscape of spliceosome factors and provides a framework for understanding the contribution of SF3B1 dysregulation to mis-splicing and splicing related disease.

## Funding

The authors are thankful for the financial support provided by MUR-PRIN2022 Project CUP B53D23025350001. This work benefited from access to the CERM/CIRMMP research Infrastructure, part of the INSTRUCT-ERIC research infrastructure. Financial support was provided by the European Union’s Horizon Europe research and innovation programme under grant agreement number 101058620 (canSERV, PID 39513). A.M. acknowledges the CINECA award under the ISCRA initiative for providing high-performance computing resources and support. A.M. and S.O. thanks the Italian Association for Cancer Research (Project AIRC IG 24514, AIRC IG 20778)). R.P. acknowledges support of the NextGenerationEU programme within the framework of the NRRP project CN00000041 (CUP B13C22001010001) and project IR0000009 (CUP B53C22001790006) as well as the HORIZON-WIDERA project MR LATVIA (no. 101160091). S.D.M. acknowledges the Italian National Centre for HPC, Big Data, and Quantum Computing (Grant No. CN00000013), founded within the Next Generation EU initiative.

## Supporting information

Supplemetari tables S1-S3, Suppementary Figures S1-S21

## Acknowledgements

The authors thank Dr. D. Blazek for providing the SF3B1(1–463) plasmid.

## Author Contributions

S.O., K. B, M.L.R.N., R.M., M.G., S. D.M., T. S., and B. P.O. S. contributed to Investigation, Data curation, Formal analysis, Validation, Visualization, L.G., S.O., I.C. F.., R. P., and A.M. contributed to Conceptualization and Methodology. S.O., K.B, M. L R N., I.C. F., R.P and A.M Writing – original draft. R.P. and A.M. were responsible for Funding acquisition, Project administration, Resources, and Supervision. All authors contributed to Writing – review and editing, interpretation of the results, and approved the final version of the manuscript.

## Declaration of Competing Interest

The authors declare no competing financial interest

## Data availability

Computational data that support the findings of this study are openly available in Zenodo https://doi.org/10.5281/zenodo.23133036, NMR data have been deposited in the Biological Magnetic Resonance Data Bank (BMRB) under accession number 54062, SAXS data are available at the following link https://www.sasbdb.org/project/2996/h64q8bjlrp/

## Notes

https://doi.org/10.5281/zenodo.23133036,

https://www.sasbdb.org/project/2996/h64q8bjlrp/

### Competing Interest Statement

The authors have declared no competing interest.

