## Supplementary material for "Charting the Structure and Functional Landscape of the SF3B1 Intrinsically Disordered Region": Supplemetari tables S1-S3, Suppementary Figures S1-S21

**Content**

Supplementary Table S1-S2 Page 2-3

Supplementary Figure S1-S21 Page 4-21

### Supplementary Tables

**Table S1.** Nucleotide sequences used in this study.

| Name | nucleotide sequence (5'-3') |
|---|---|
| ***ssDNA 20nt*** | **[6-FAM]-**CTACTACCCCCACCCTCAC |
| ***D1*** | **[6-FAM]**-CTACTACCCCCACCCTCACAACCTTTTTTTTTTTTTTT |
| ***D3*** | TTTTTTTTTTTTTTTTGGTTGTGAGGGTGGGGGTAGTAG |
| ***ssRNA 12 nt*** | **[6-FAM]**- cuacuaccccca |
| ***ssRNA 20nt*** | **[6-FAM]**- cuacuacccccacccucaca |
| ***ssRNA 27nt*** | [**6-FAM**]-cuacccccacccucacaaccuuuuuuuuuu |
| ***R3*** | uuuuuuuuuuuuuuuugguugugaggguggggua |
| ***D11*** | GTTGTAAAACGACGGCCAGTGCCTTTTCCCAGCCTCAATCTCATCACTCTAG AGGATCCCCGGGTAC |
| ***D9*** | **[6-FAM]-** GTACCCGGGGATCCTCTAGAGTCGACCTGCAGGCATGCAAGCTTGGCACTGG CC GTCGTTTTACAAC |
| ***D4*** | GCTTGCATGCCTGCAGGCCAGCCTCAATCTCATC |
| ***R4*** | gcuugcaugccugcaggccagccucaaucucaucuuuuu |

**Table S2.** SAXS parameters of the analyzed proteins.

| | RG (nm) | Dmax (nm) |
|---|---|---|
| $^{186\text{–}341}$SF3B1 | 3.6 +/- 0.1 | 10 +/- 1 |
| GST-$^{1\text{-}463}$SF3B1 | 6.57 +/- 0.2 | 20 +/- 2 |

**Table S3**: Details of molecular dynamics simulations. The first column reports the simulated system in the wild-type (WT) or phosphorylated (4p) state; the second column reports the force field and the water model employed in MD simulations. The third and fourth column report the simulation lengths and the number of replicas.

| **SF3B1 System** | **Force Field + Water Model** | **Time (μs)** | **Number of replicas** |
|---|---|---|---|
| SF3B1(186-341) WT | C36m + TIP4PD | 2 | 3 |
| SF3B1(186-341) 4p | | 2 | 3 |
| SF3B1(186-341) WT | A99SB-ILDN + TIP4PD | 2 | 1 |
| SF3B1(186-341) WT | A99SB-DISP + DISP | 2 | 3 |
| SF3B1(186-341) 4p | | 2 | 3 |

### Supplementary Figures

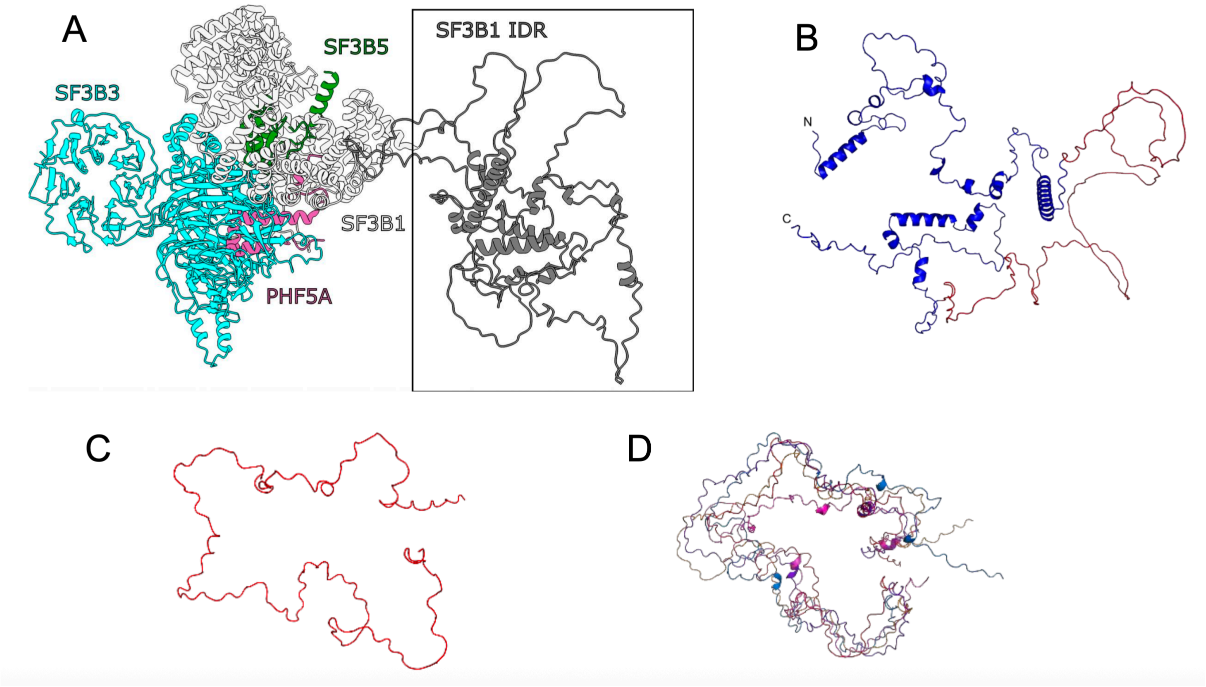


**Figure S1.** AlphaFold3 predicted models of SF3b and SF3B1 constructs. (A) Structure of the SF3b complex. SF3B3, SF3B5, PHF5A, and SF3B1 are shown in blue, green, pink, and white cartoon representations, respectively. The intrinsically disordered N-terminal region of SF3B1 is displayed on the right. (B) Full NTD (residues 1-463) SF3B1 fragment showing a largely disordered structure, with several regions predicted to adopt short α -helical elements. The 186-341 SF3B1 segment is highlighted in red, illustrating that this portion remains completely disordered within the full NTD. (C) SF3B1 construct (residues 186-341, red), modelled as entirely disordered and (D) overlay of five predicted conformers of the 186-341 SF3B1 construct, illustrating the intrinsic disorder and conformational variability of the segment.

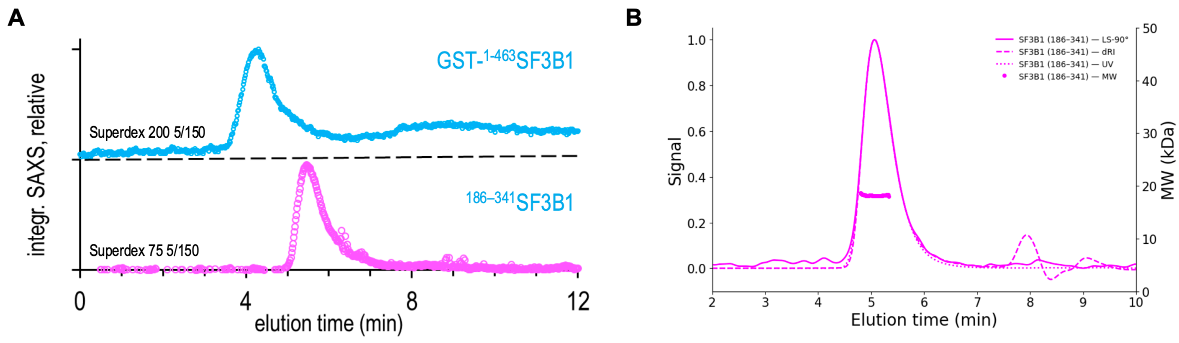


**Figure S2.** SEC–UV/SAXS analysis of the peptide and GST fusion protein. Samples were injected onto a Superdex 75 5/150 size-exclusion chromatography column at a flow rate of 0.3 mL min$^{-1}$. During elution, SAXS data were collected in 0.5 s frames for the peptide and 1 s frames for the GST fusion protein. (A) UV absorbance recorded at 280 nm, shown as the elution profile versus elution time (min). (B) Integrated SAXS intensity profiles generated using Chromixs.

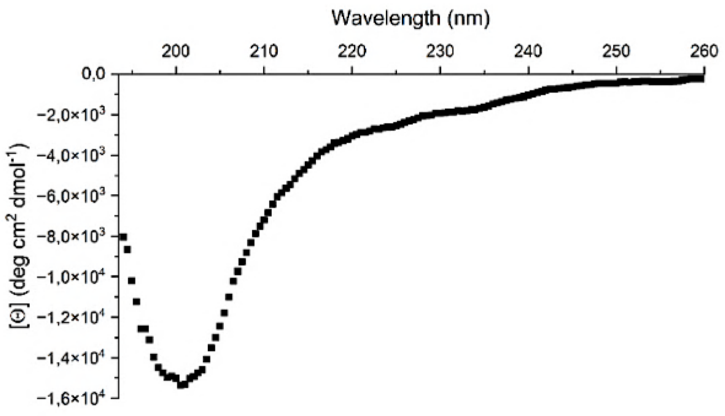


**Figure S3.** Far-UV CD spectrum of the $^{186-341}$SF3B1 fragment. Molar ellipticity [Θ] is plotted as a function of wavelength (180-260 nm). The spectrum exhibits a single minimum near 200 nm, indicative of a predominantly random coil conformation and consistent with the protein intrinsically disordered nature.

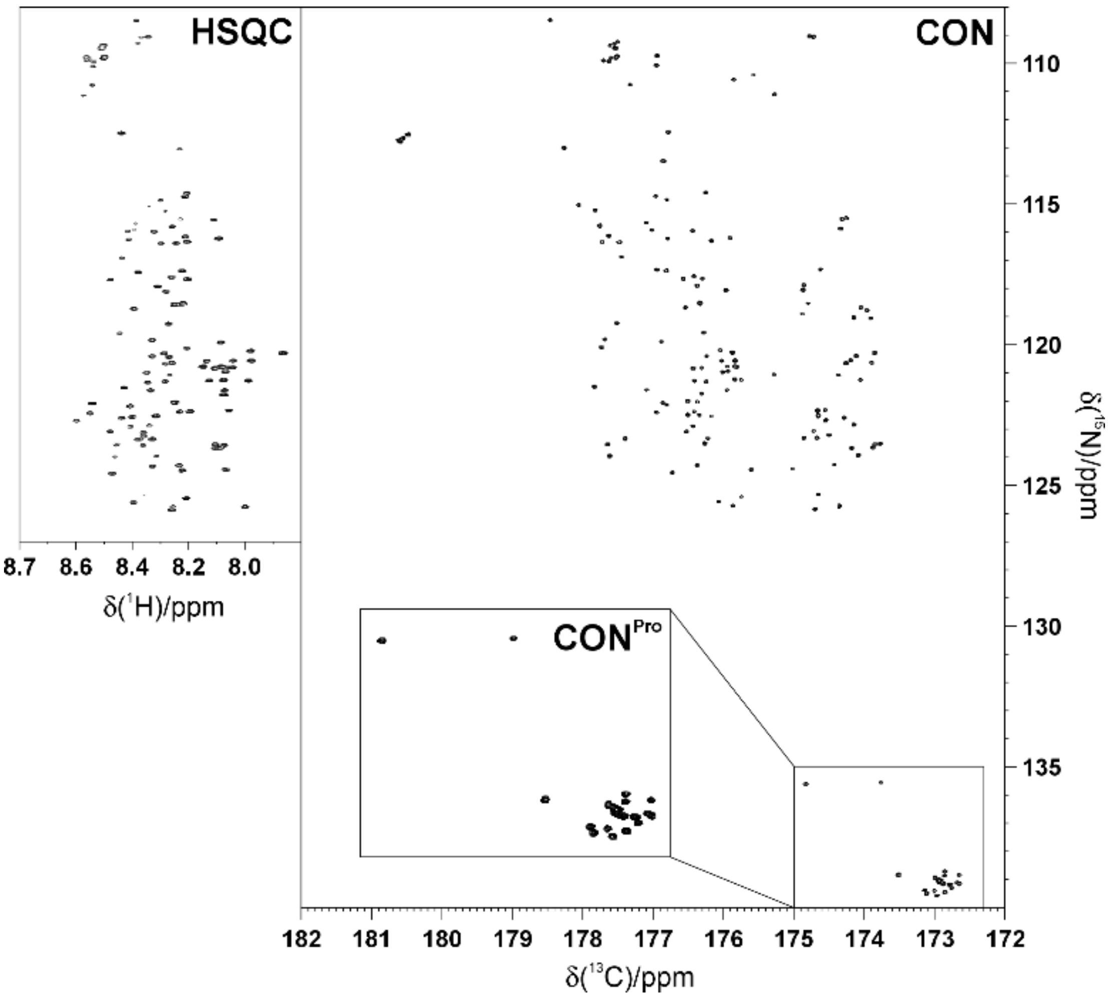


**Figure S4.** $^{1}H^{15}N$-HSQC and CON spectra of $^{186-341}$SF3B1 construct recorded at 298 K and 28.2 T (1200 MHz). The HSQC spectrum illustrates the limited $^{1}H$ chemical shift dispersion typical of disordered proteins. The corresponding CON spectrum is shown together with a zoomed view of the proline region, where signals from the TP repeats cluster in a narrow spectral window. The zoom highlights 17 resonances with highly similar $^{13}C$ and $^{15}N$ chemical shifts.

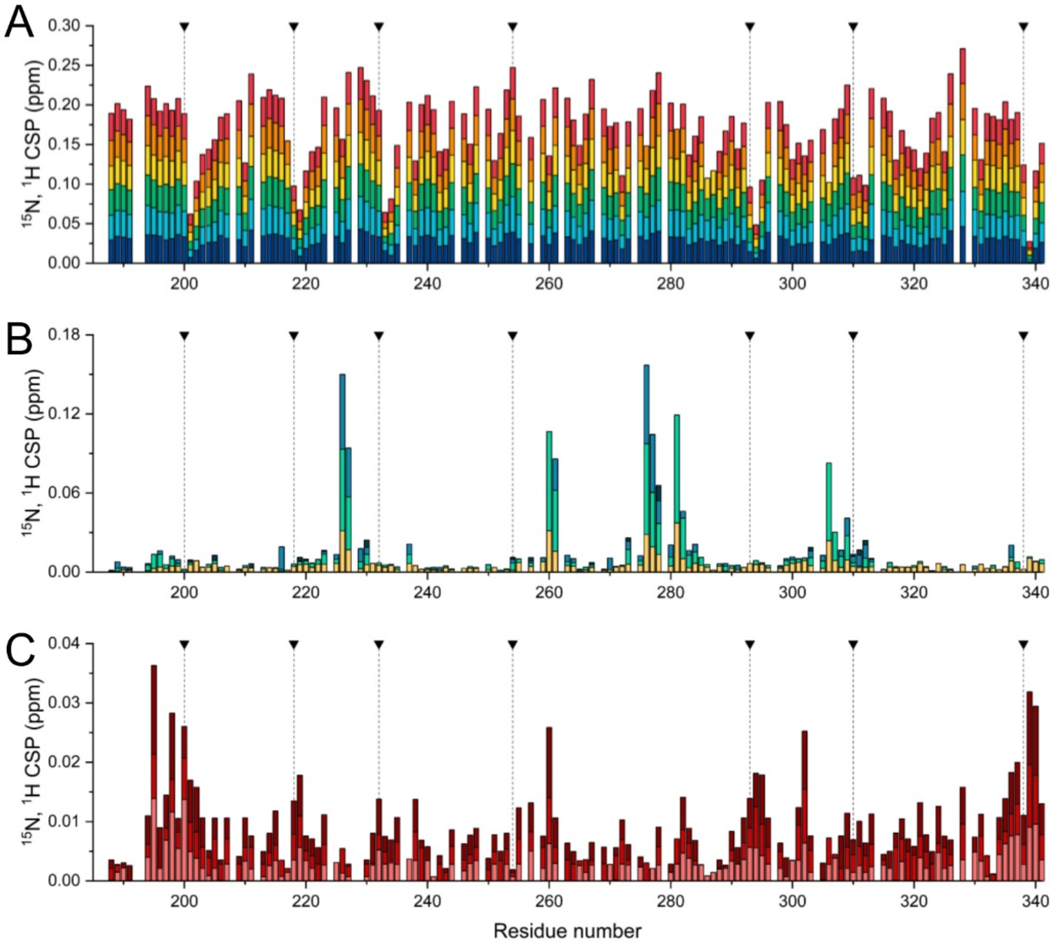


**Figure S5.** Residue-specific responses of $^{186\text{–}341}$SF3B1 to temperature, pH, and ionic strength. (A) Cumulative $^{1}$H and $^{15}$N chemical shift perturbation (CSP) upon temperature changes, calculated relative to 278 K and colour-coded from blue (smallest shift, 283-278 K) to red (largest shift, 308-278 K). (B) CSP as a function of pH, calculated relative to pH 5.5 and colour-coded from yellow (pH 6.0) to dark blue (pH 8.0). (C) CSP as a function of ionic strength (NaCl concentration), calculated relative to 0 mM NaCl and colour-coded from light red (50 mM NaCl) to dark red (200 mM NaCl). Black triangles with dashed vertical lines indicate the positions of tryptophan residues.

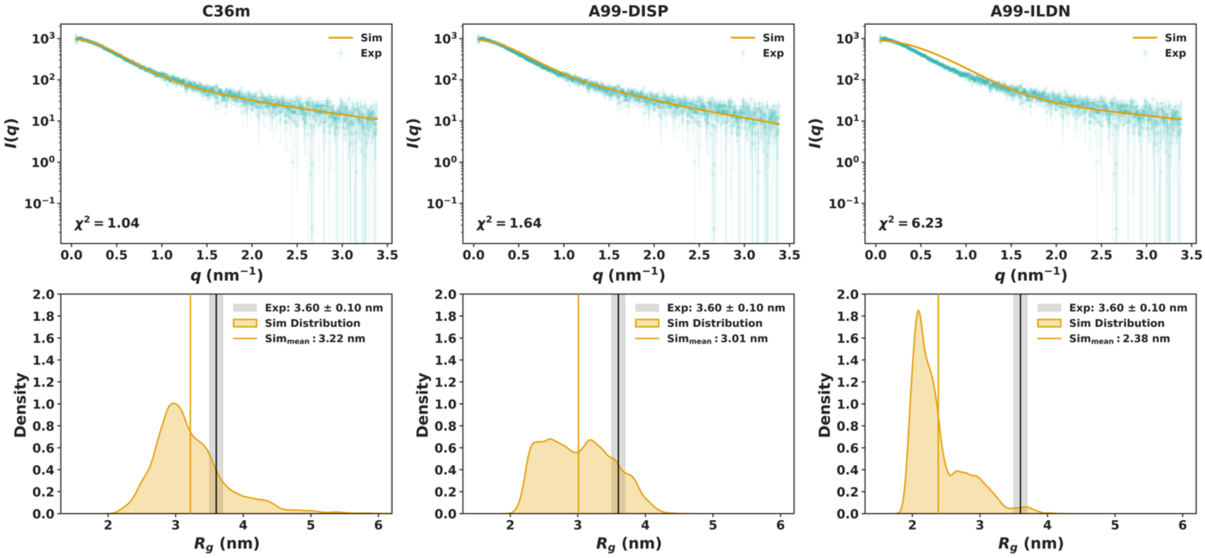


**Figure S6.** Force-field-dependent agreement of $^{186–341}$SF3B1 conformational ensembles with SAXS data. Top panels: Experimental SAXS profiles (green) compared with scattering curves calculated from molecular dynamics ensembles generated using the C36m, A99-DISP, and A99-ILDN force fields (orange). The quality of the agreement is quantified by the corresponding $\chi^2$ values. Bottom panels: Distributions of the radius of gyration ($R_g$) sampled by each ensemble. The vertical grey line and shaded region indicate the SAXS-derived experimental $R_g$ (3.60±0.10 nm), while the orange line denotes the mean simulated Rg.

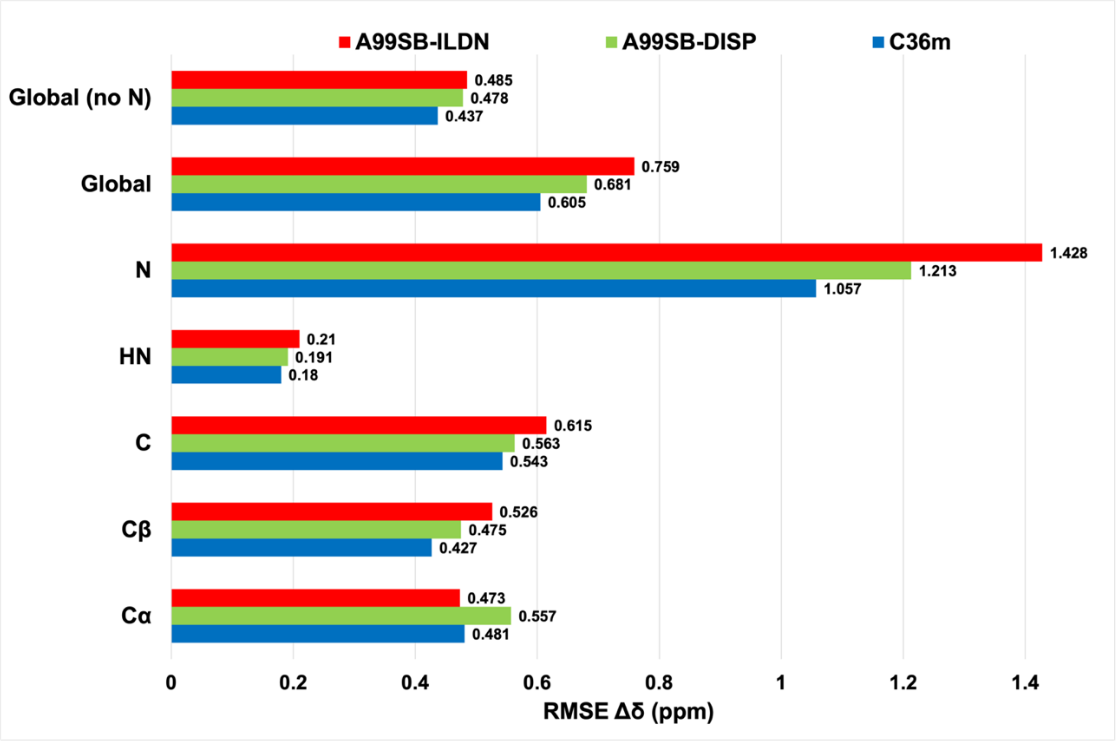


**Figure S7.** Comparison of simulated and experimental NMR chemical shifts for the wild-type $^{186–341}$SF3B1 fragment. Root Mean square error (RMSE) is computed for the different types of nuclei and the different force fields used on MD simulation trajectory frames. Values relative Amber99-ILDN (A99-ILDN), Amber99-DISP (A99-DISP), and CHARMM-C36m (C36m) force fields are shown in red, green and blue, respectively. Amide-nitrogen ($^{15}$N) shifts were excluded from the global RMSE since they are particularly sensitive to long-range electrostatics, hydrogen bonding networks, and inconsistencies across experiments. Nonetheless, the $^{15}$N RMSE values were analysed separately, confirming that the overall trends remain consistent across nuclei.

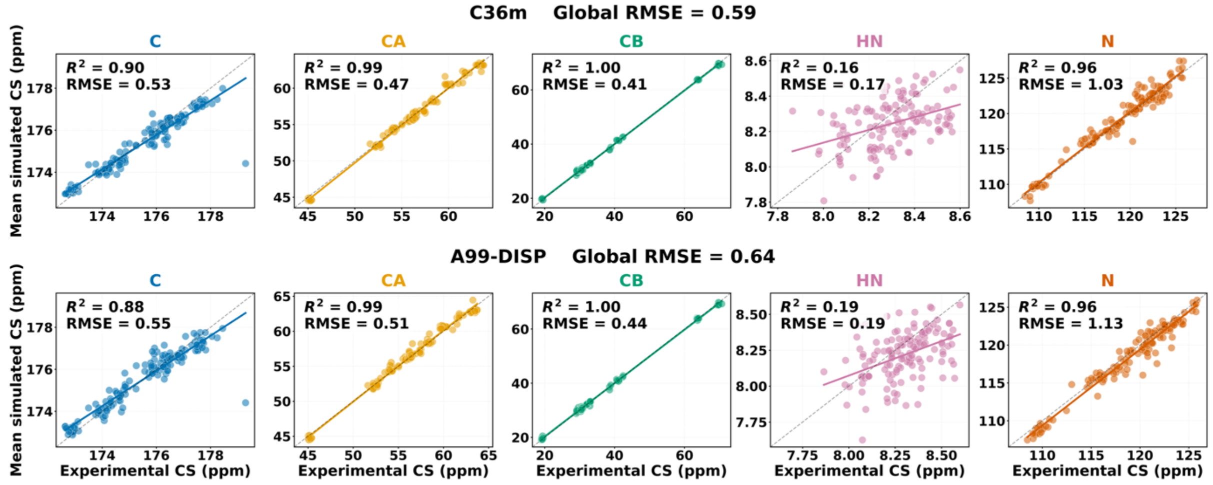


**Figure S8.** Correlation between experimental and computed NMR chemical shifts for the wild type $^{186–341}$SF3B1 fragment. The NMR chemical shifts are computed on the combined trajectories derived from 3 molecular dynamics simulation replica performed with the CHARMM36m (C36m, Top) and Amber99-DISP (A99-DISP, bottom) force field, respectively. From left to right comparison is done for C, $C^{\alpha}$, $C^{\beta}$, N and $H^{N}$ nuclei. Top and bottom panels report results obtained for NMR chemical shifts computed on the MD simulations trajectories generated with the the global root mean square error (RMSE) for C36m and A99-DISP is of 0.59 and 0.64, respectively.

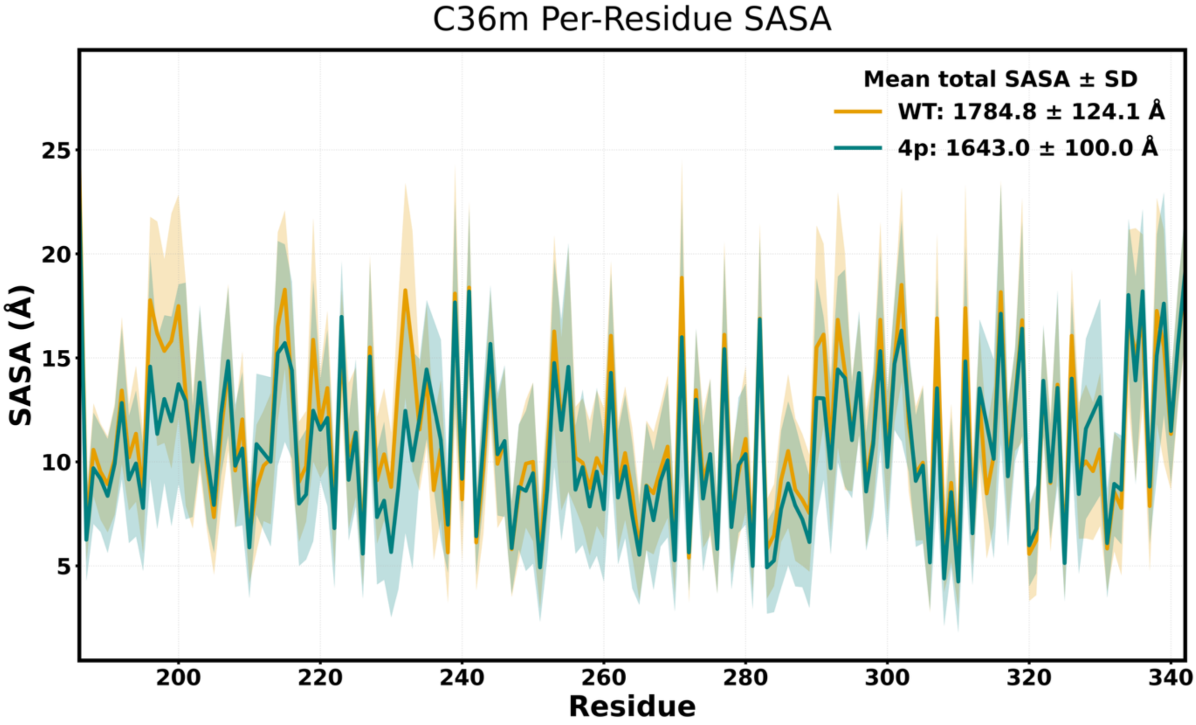


**Figure S9.** The per-residue Solvent Accessible Surface Area (SASA) computed for the wild-type (orange), and phosphorylated (green) $^{186–341}$SF3B1 fragment computed with the CHARMM-C36m (C36m) (top) force field. Shaded area indicates standard deviation.

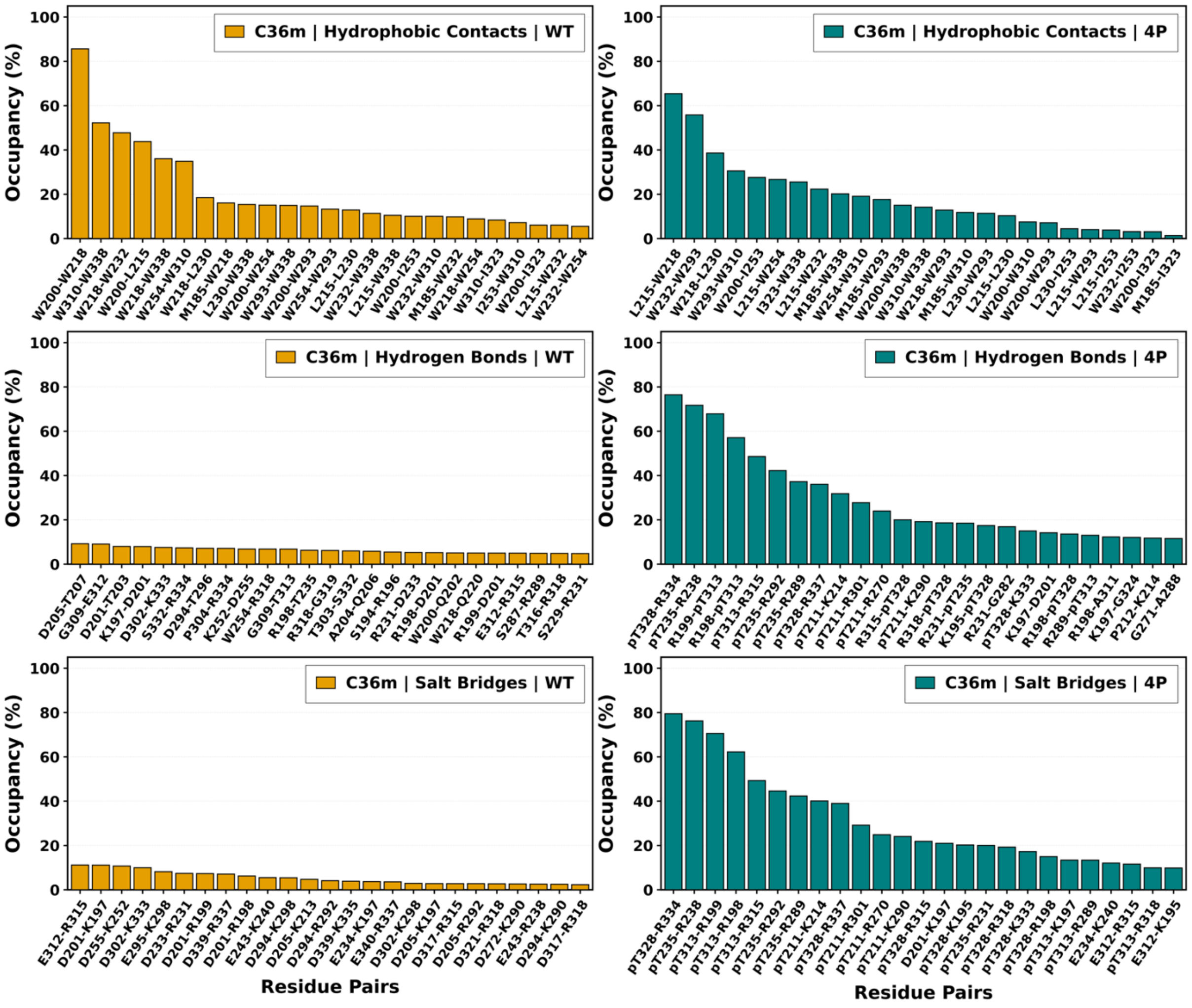


**Figure S10.** Analysis of the 25 most persistent hydrophobic, hydrogen bond, and salt bridge occupancies (%) in the wild-type and phosphorylated (pThr211, 235, 313, 328) $^{186–341}$SF3B1 fragment computed on the MD simulation trajectories with the CHARMM36m force field.

**A**

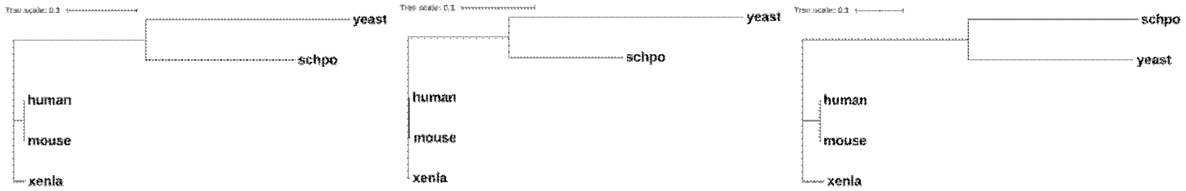


**B**

```
human           MAKIAKTHEDIEAQIREIQGKKAALDEAQGVGLDSTGYYDQEIYGGSDSRFAGYVTSIAA
yeast           -----------------------------------------------MSHPIQFV-----
                                                                *.   :*

human           TELEDDDDDYSSSTSLLGQKKPGYHAPVALLNDIPQSTEQYDPFAEHRPPKIADREDEYK
yeast           -----NANNSDKSHQLGGQ----YSIPQDLRENLQKEA-----------ARIGENEK---
                     : :: ..* .* **    *  *  * ::: :.:           ..*.:.*.

human           KHRRTMIISPERLDPFADGGKTPDPKMNARTYMDVMREQHLTKEEREIRQQLAEKAKAGE
yeast           ---------------------------------DVLQEKMETR-----------------
                                                 **:.*:  *.

human           LKVVNGAAASQPPSKRKRRWDQTADQTPGATPKKLSSWDQAETPGHTPSLRWDETPGRAK
yeast           ------TVQNREDSYHKRRFDM--------------------------------------
                      :. ..  * .***:*

human           GSETPGATPGSKIWDPTPSHTPAGAATPGRGDTPGHATPGHGGATSSARKNRWDETPKTE
yeast           ---------------------------------------------------KFEPDSDTQ
                                                                   .::  ..*:

human           RDTPGHGSGWAETPRTDRGGDSIGETPTPGASKRKSRWDETPASQMGGSTPVLTPGKTPI
yeast           TVTSSENTQDAVVPR-----------------KRKSRWD---------------------
                  *.. .:  * .**                 *******

human           GTPAMNMATPTPGHIMSMTPEQLQAWRWEREIDERNRPLSDEELDAMFPEGYKVLPPPAG
yeast           ------------------------------------------------VKGYE--PPDES
                                                                 :**:  **  .
```

**Figure S11.** (A) Phylogenetic trees of orthologous proteins, shown from left to right: full-length protein, C-terminal domain (CTD), and N-terminal domain (NTD). While the CTD is highly conserved across vertebrates, the NTD exhibits greater variability, particularly between vertebrates and fungi, consistent with its intrinsically disordered nature. (B) MUSCLE alignment of the human and yeast N-terminal regions. Conserved regions are highlighted in red, and the human fragment analysed (residues 186-341) shown in bold.

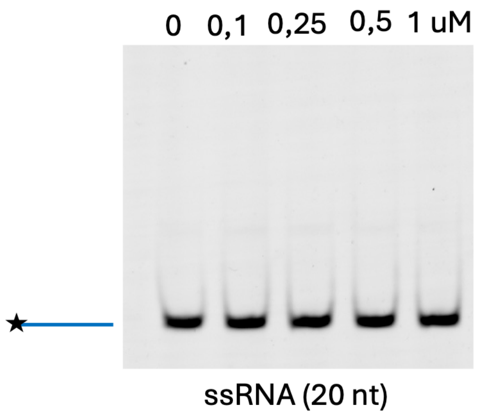


**Figure S12:** Electrophoretic mobility shift assay (EMSA) showing the binding of GST to a 20-nt ssRNA substrate. GST was incubated with increasing concentrations of protein (0, 0.1, 0.25, 0.5 and 1 µM) in the presence of 10 nM fluorescently labelled ssRNA. No detectable shift in RNA mobility was observed, indicating that GST alone does not bind ssRNA under the experimental conditions used.

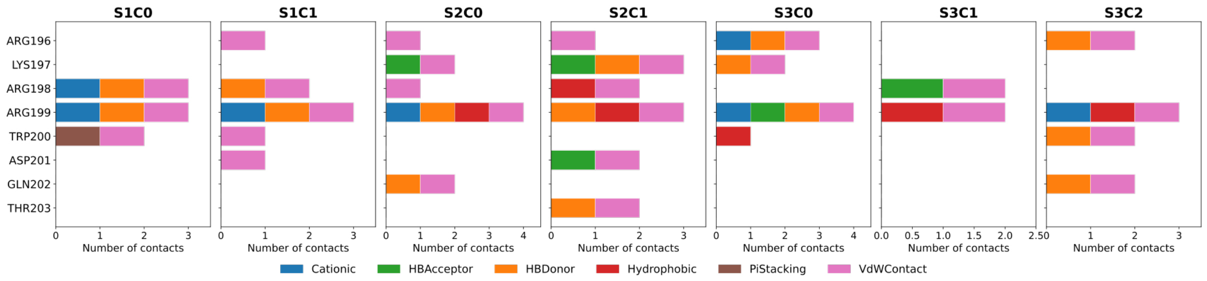


**Figure S13.** Interactions of the W200-centered SF3B1 fragment (i.e. fragment RKRRW[200]DQTA) for the three RNA splits. Naming refers to split number and cluster number (S1C0 stands for RNA split1 cluster 0). Interactions are listed as Anionic' RNA binding via the phosphate groups (brown); HBAcceptor (blue) means that the RNA is the hydrogen bond acceptor. HBDonor (red) means that the RNA is the hydrogen bond donor, hydrophobic (green), π–π-staking (violet) and van der Waals (gray) interactions are also reported.

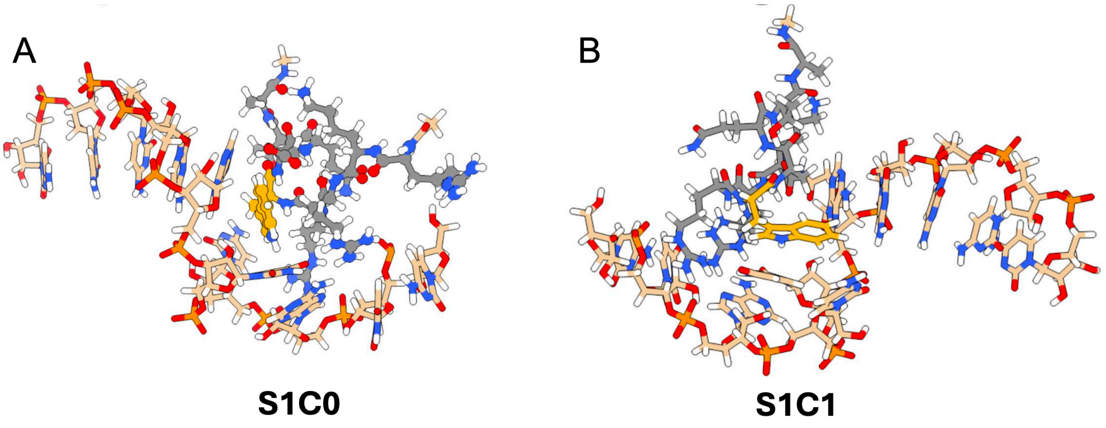


**Figure S14.** AlphaFold3-generated starting structures of RNA split 1 in complex with the W200-containing SF3B1 fragment (RKRRW$^{200}$DQTA). Panel A shows the initial structure corresponding to cluster 0, while panel B corresponds to cluster 1. Carbon atoms of SF3B1 and RNA are displayed in grey and pink, respectively, with W200 highlighted in orange. Nitrogen, oxygen, phosphate, and hydrogen atoms are shown in blue, red, gold, and white, respectively.

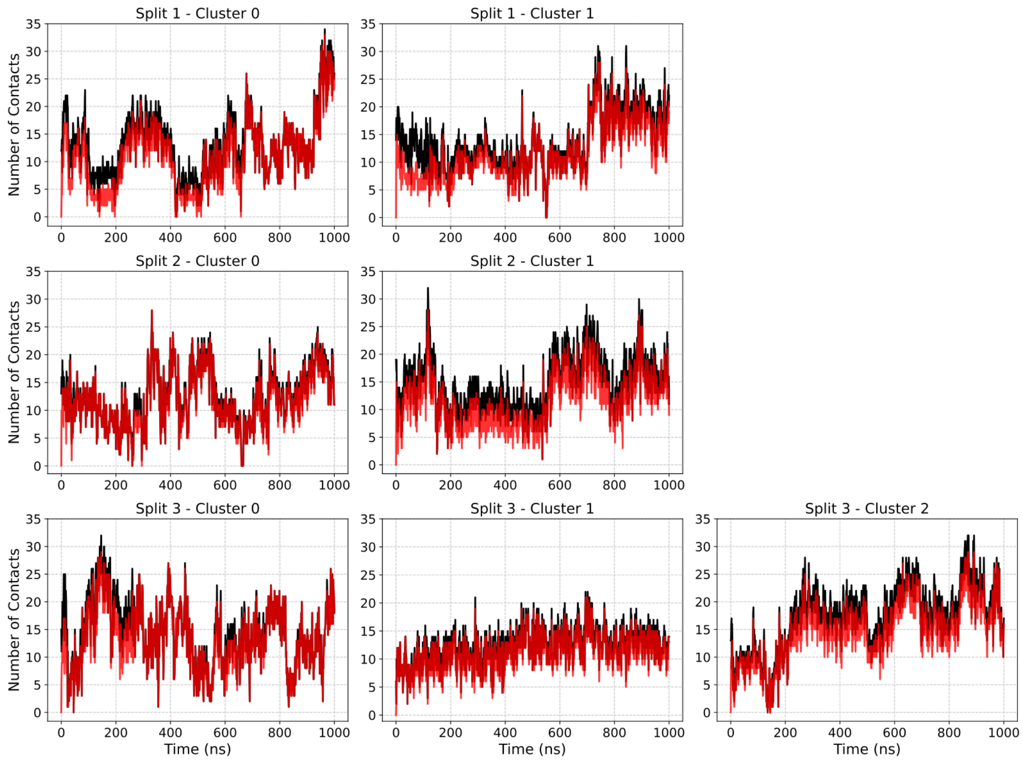


**Figure S15.** Total (black) and newly (red) formed contacts as a function of molecular dynamics simulation time (ns) of Alpha Fold3-generated adducts between the three RNA segments and the W200-containing SF3B1 fragment. Panels within each row represent different initial conformations of each adduct.

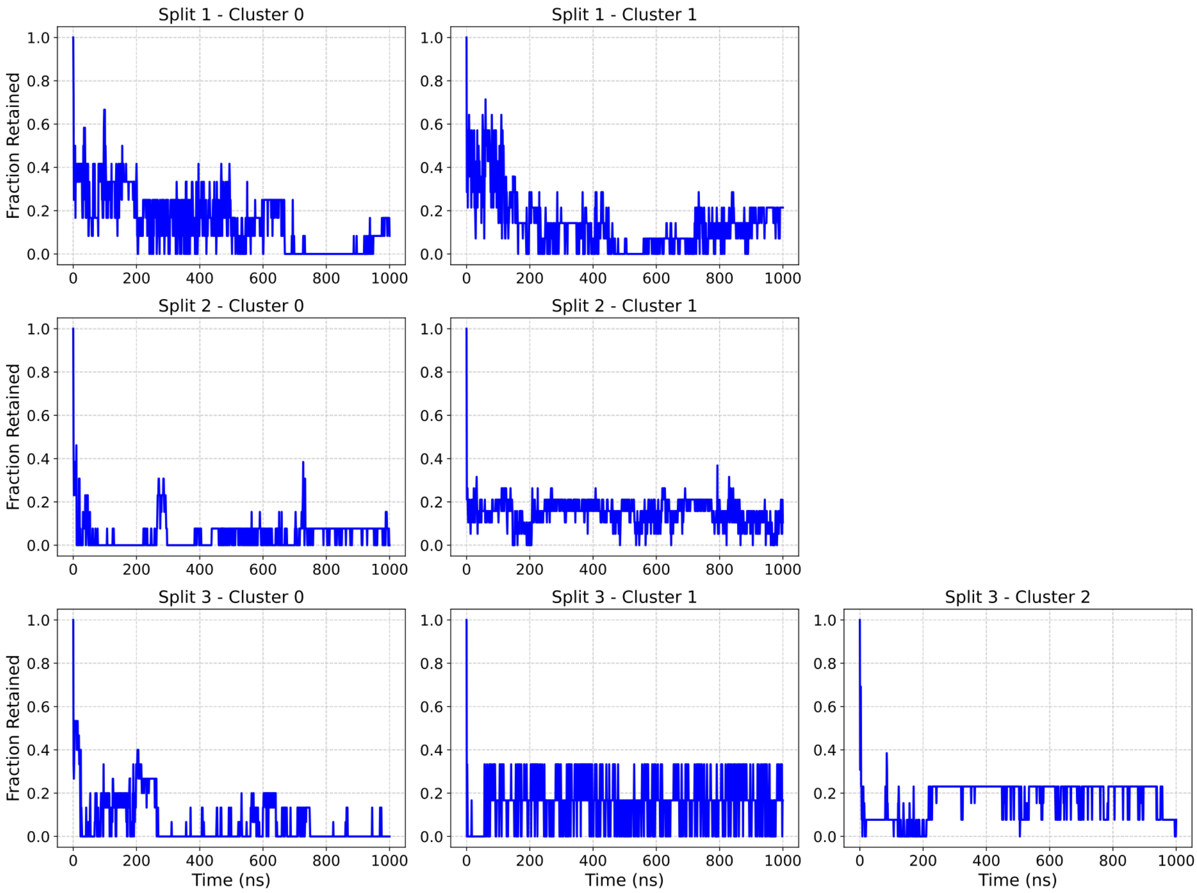


**Figure S16.** Fraction of native contacts retained as a function of molecular dynamics simulation time (ns) of AlphaFold3-generated adducts between the three RNA segments and the W200-containing SF3B1 fragment. Panels in each line correspond to different initial conformations for each adduct.

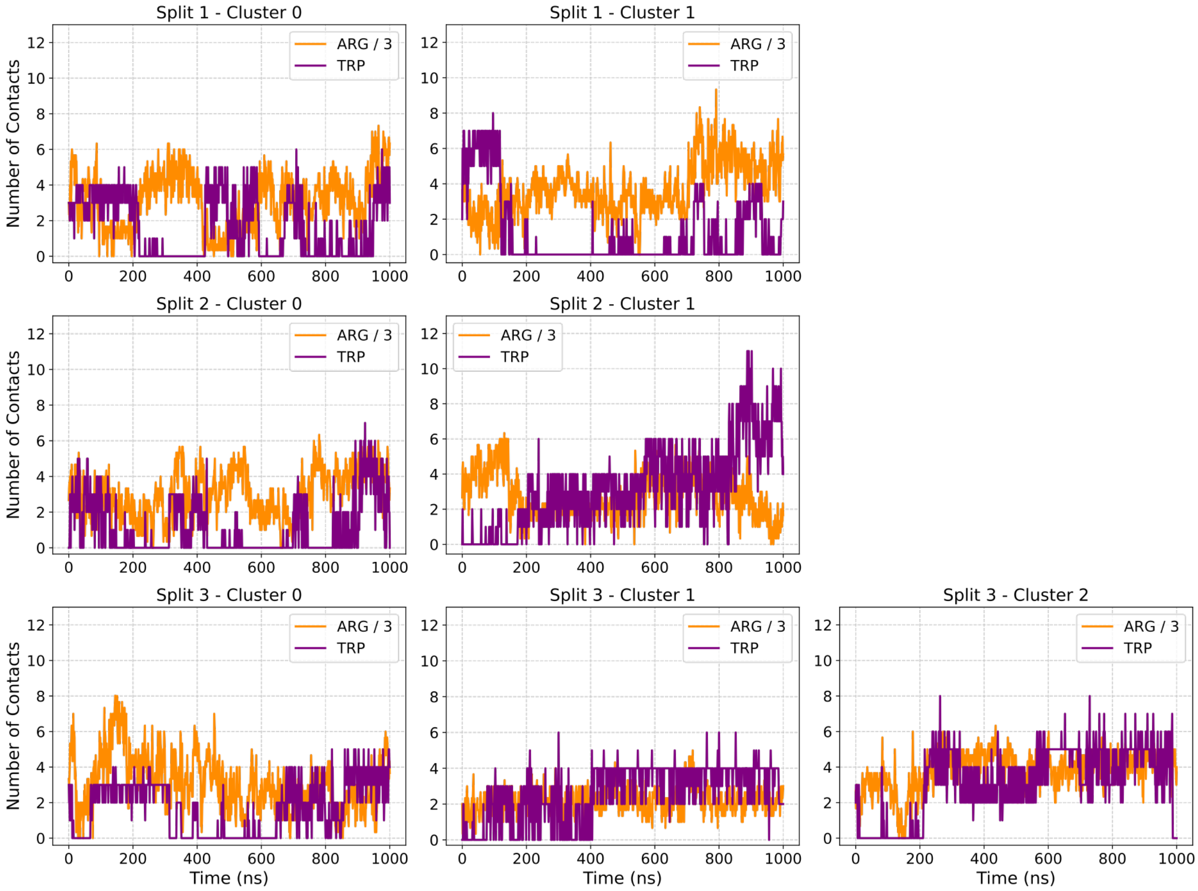


**Figure S17.** Normalized number of contacts formed by R196, R198, R199 (orange), and W200 (purple) with the different RNA segments as a function of molecular dynamics simulation time (ns) of AlphaFold 3-generated adducts between the three RNA splits and the W200-containing SF3B1 fragment. Panels within each row correspond to distinct initial conformations for each adduct.

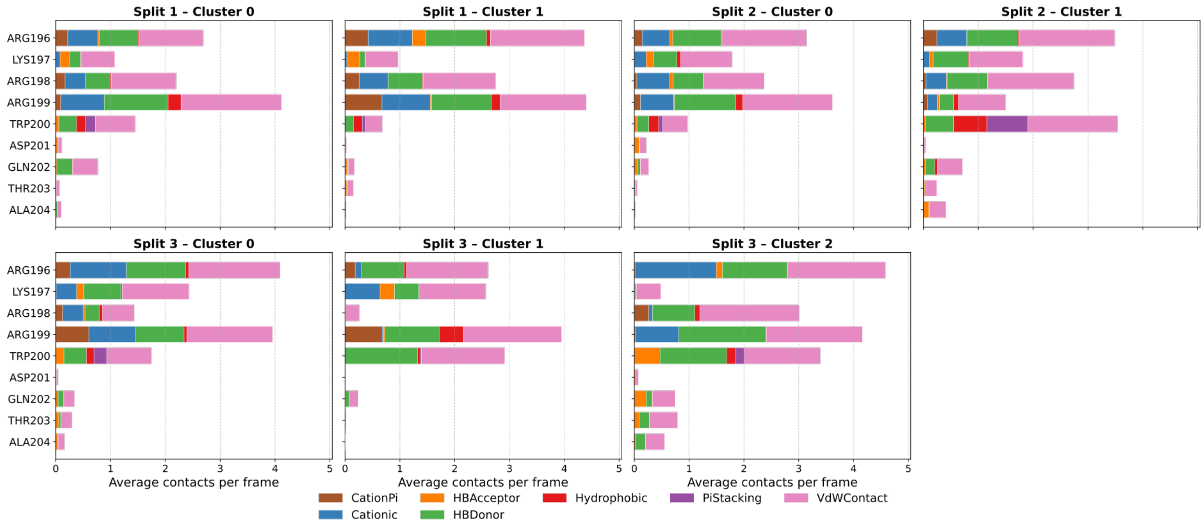


**Figure S18.** Per-residue interaction profile between the W200-centered SF3B1 fragment (RKRRW$^{200}$DQTA) and the three RNA segments. Interactions are categorised as anionic (RNA phosphate-mediated; brown), hydrogen bond acceptor (RNA acting as hydrogen bond acceptor; orange), hydrogen bond donor (RNA acting as hydrogen bond donor; blue), hydrophobic (green), π–π stacking (violet) and van der Waals (grey).

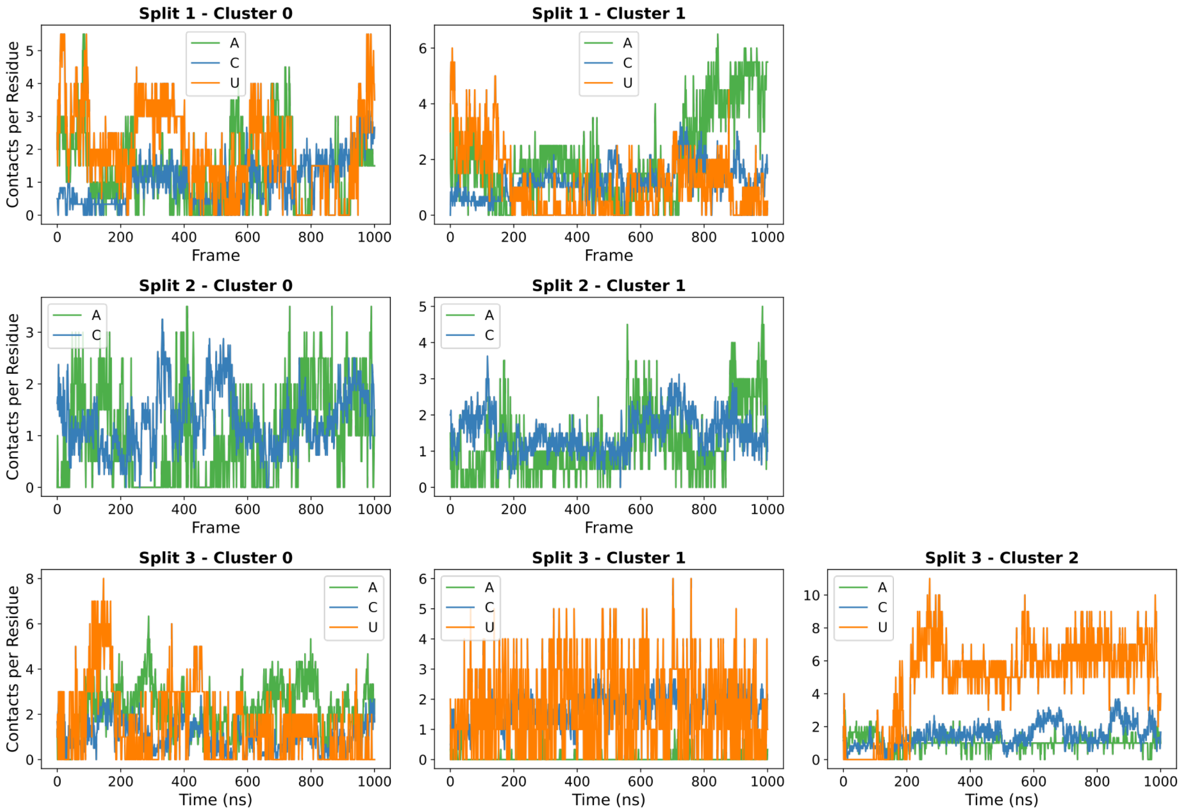


**Figure S19.** Normalized contact frequencies per nucleobase as a function of molecular dynamics simulation time (ns) of the W200-centered SF3B1 fragment (RKRRW$^{200}$DQTA) in complex with the three RNA segments. Interactions involving A, U, and C are shown in blue, orange, and green, respectively.

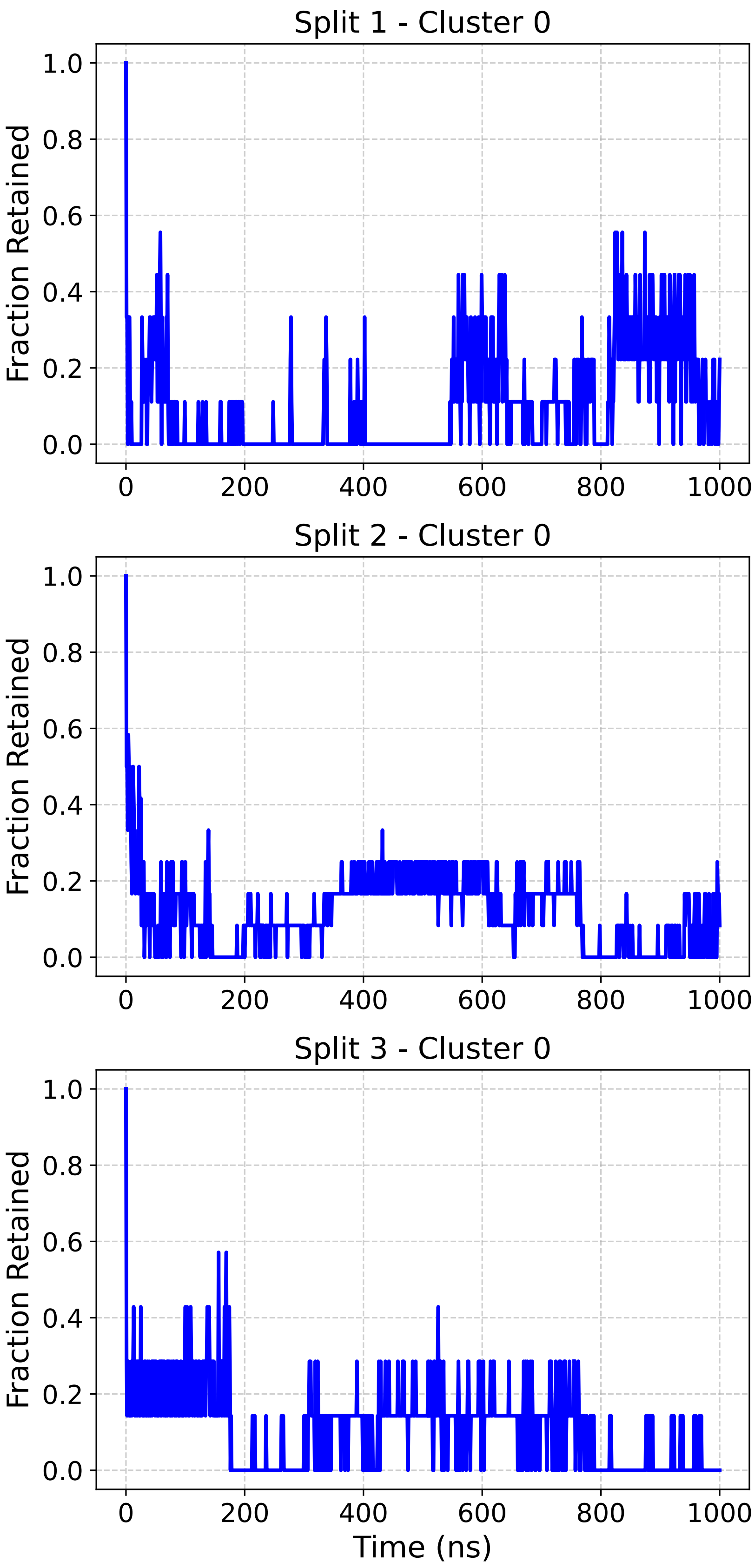


**Figure S20.** Fraction of native contacts retained as a function of molecular dynamics simulation time (ns) of AlphaFold3-generated adducts between the three RNA segments and the K240-containing SF3B1 fragment. Panels in each line correspond to a different RNA split.

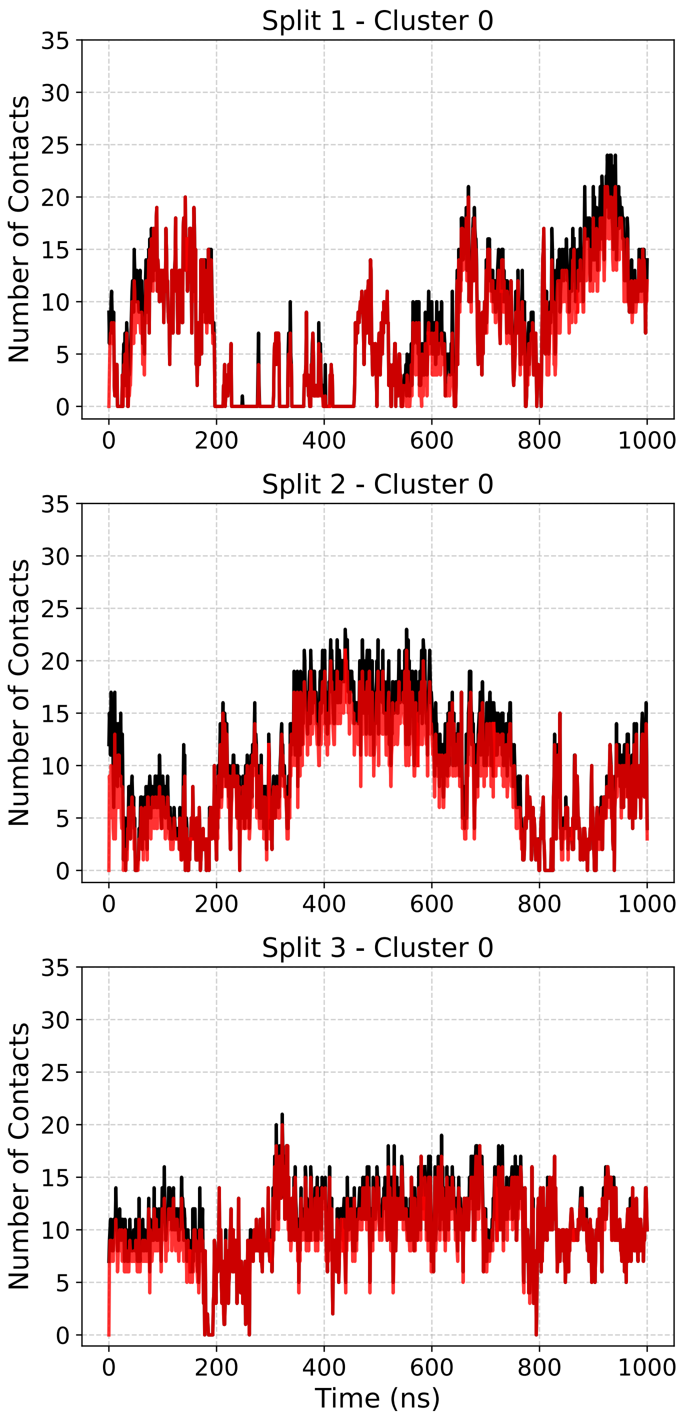


**Figure S21.** Total (black) and newly formed (red) contacts as a function of molecular dynamics simulation time (ns) of AlphaFold3-generated adducts between the three RNA segments and the K240-containing SF3B1 fragment. Panels within each row represent a different RNA split.
